# TFEB controls syncytiotrophoblast differentiation

**DOI:** 10.1101/2024.02.20.581304

**Authors:** Meagan N. Esbin, Liza Dahal, Vinson B. Fan, Joey McKenna, Eric Yin, Xavier Darzacq, Robert Tjian

**Affiliations:** Department of Molecular and Cell Biology, University of California Berkeley, Berkeley, CA; Howard Hughes Medical Institute, University of California, Berkeley, CA

## Abstract

During human development, a subset of differentiating fetal cells form a temporary organ, the placenta, which invades the uterine wall to support nutrient, oxygen, and waste exchange between the mother and fetus until birth. Most of the human placenta is formed by a syncytial villous structure which arises via cell-cell fusion of underlying fetal trophoblast stem cells. Genetic and functional studies have characterized the membrane protein fusogens, Syncytin-1 and Syncytin-2, that are both necessary and sufficient for human trophoblast cell-cell fusion. However, identification and characterization of upstream transcriptional regulators regulating their expression has been limited. Here, using CRISPR knockout in an *in vitro* cellular model of syncytiotrophoblast development (BeWo cells), we find that the transcription factor TFEB, mainly known as a regulator of autophagy and lysosomal biogenesis, is required for cell-cell fusion of syncytiotrophoblasts. TFEB translocates to the nucleus, exhibits increased chromatin interactions, and directly binds the Syncytin-1 and Syncytin-2 promoters to control their expression during differentiation. While TFEB appears to play an important role in syncytiotrophoblast differentiation, ablation of TFEB largely does not affect lysosomal gene expression or lysosomal biogenesis in differentiating BeWo cells, suggesting that TFEB plays an alternative role in placental cells.

## Introduction

Shortly after implantation of a fertilized blastocyst, the placenta, a temporary organ, begins to develop alongside the human fetus which provides vital sustenance and exchange during pregnancy and sets the fetus up for a healthy life post-birth.^1–3^ Many of the underlying molecular mechanisms driving development of the placenta remain enigmatic yet vital to understand. The placenta is thought to be the basis of many obstetrical complications, so its dual role in maintaining both fetal and maternal health during pregnancy suggests that a deeper understanding of human placental development provides an opportunity to both support healthy neonate development and combat the rising maternal mortality rates plaguing the U.S and the world.^4–6^ During early development of the human placenta, trophoblast stem cells generate three main lineages where cytotrophoblast progenitor cells (CTB; a mononuclear stem cell which proliferates and supports placental growth) differentiate into two main functional cell types: syncytiotrophoblasts (STB; a multinucleated syncytium formed via cell-cell fusion of cytotrophoblasts that forms the outermost barrier between the fetal and maternal circulation), and extravillous trophoblasts (EVT; a mononuclear specialized subset of cytotrophoblasts which invade and remodel the maternal spiral arteries to allow vascularization of the placenta).^7^ The regulation of syncytialization within the placental villous appears vital for placental and maternal health. Aberrant syncytialization has been observed concomitant with several important obstetrical conditions including in preeclampsia (reduced syncytialization) and in fetal growth restriction (excess syncytialization).^8–10^

Differentiation of syncytiotrophoblasts is characterized by cell-cell fusion, secretion of pregnancy hormones including hCG, nuclear enlargement, and exit from mitosis while differentiated cells remain transcriptionally active.^11–13^ The molecular mediators of cell-cell fusion in human syncytiotrophoblasts are evolutionarily modern endogenous retroviral envelope proteins, Syncytin-1 and Syncytin-2, which are membrane proteins with high expression in the cytotrophoblast and syncytiotrophoblasts of the human placenta.^14–17^ Syncytin-1 and -2 have fusogenic activity and are sufficient to induce cell-cell fusion in cells expressing their receptors, SLC1A4/5 and MFSD2A, respectively.^15,18,19^ Decades of research have illustrated the important role of many cellular factors in orchestrating the differentiation and cell-cell fusion of syncytiotrophoblasts, but only one direct upstream regulator of placental syncytialization has been identified, the transcription factor GCM1.^20^ Interestingly, GCM1 has been co-opted evolutionarily to be capable of regulating the sequence-unrelated syncytin genes in both human^21–23^ and mouse ^24–27^ and GCM1 has been shown to directly bind the promoters of Syncytin-1, Syncytin-2, and MFSD2A.^23,28,29^ To date, other direct regulators of Syncytin-1 or -2 expression have not been identified.

The transcription factor TFEB is a potentially interesting, conserved candidate for regulating placental differentiation. Like GCM1, homozygous knockout of the transcription factor *TFEB* is lethal during gestation (d9.5) due to failure to form the syncytialized labyrinth layer of the murine placenta.^27,30^ TFEB belongs to a small family (along with TFE3, MITF, and TFEC) of basic helix-loop-helix DNA-binding transcription factors that homo- and heterodimerize to bind a subset of E-box motifs in the genome.^31,32^ In human cells, TFEB has been described as a master regulator of the coordinated autophagy and lysosomal biogenesis (CLEAR) gene network.^33,34^ TFEB’s transcriptional role is potentiated upon cytoplasmic-to-nuclear import which is negatively controlled largely via phosphorylation of TFEB by mTOR and other kinases.^35–37^

In patients with the placental disease preeclampsia, protein levels of TFEB along with lysosomal proteins LAMP1 and LAMP2 are reduced in placental tissue compared to age-matched controls.^38,39^ Indeed, perturbation of mTOR signaling has been shown to affect trophoblast syncytialization: in BeWo cells, treatment with the mTOR inhibitor Rapamycin increases cell fusion and hCG production, while concomitant treatment with the mTOR activator MHY1485 erases these effects.^40^ In contrast, for cases of fetal growth restriction (FGR), researchers have found reduced mTOR phosphorylation and excess cell-cell fusion.^41^ While these findings may point to TFEB as an interesting potential candidate in syncytiotrophoblast differentiation, to date TFEB’s direct role in human placental development has not been investigated.

## Results

To assess TFEB’s roles in human syncytiotrophoblast differentiation, we used CRISPR Cas9 to genetically delete TFEB in BeWo cells, an in vitro model of syncytiotrophoblast differentiation and which inducibly differentiate and fuse upon treatment with the cAMP activator Forksolin.^42–44^ To prevent genetic compensation by other TFE family members such as TFE3, we started by creating a homozygous TFEB and TFE3 double knockout BeWo cell line by transfecting plasmids encoding Cas9-Venus and 10 sgRNAs targeting the N-terminal and C-terminal regions of TFEB and TFE3 to induce genetic deletions within the two protein-coding loci (Fig. 1A, Table1). One clone (#C2) was confirmed to have deletions in the coding regions of TFEB and TFE3 and complete loss of TFEB and TFE3 protein expression by western blotting (Fig. 1B-C). While TFEB was moderately expressed in wild-type BeWo cells, TFE3 was virtually undetectable by western blotting due to its low expression (Fig. 1C, S1A). Since TFE3 was not appreciably expressed in wild-type BeWo cells we additionally created TFEB-only KO BeWo cells by nucleofecting BeWo cells with Cas9 RNPs containing a single sgRNA targeted to induce indels at the N-terminus of TFEB. The edit yielded one clone (TFEB KO #c13) with almost complete deletion of TFEB via induced indels. By western blotting, this clone had ∼11% remaining TFEB expression (Fig. 1B). Notably, the TFEB KO BeWo cells, like the wild-type, do not show any appreciable TFE3 expression indicating that TFEB’s loss is not compensated by upregulation of TFE3 in BeWo cells (Fig. 1C). In both clones, some characteristics of differentiation appear entirely unperturbed upon TFEB loss including the ability to produce hCG and the enlargement of nuclei upon differentiation with Forskolin (Fig. 1D, S1B).

**Figure 1.**
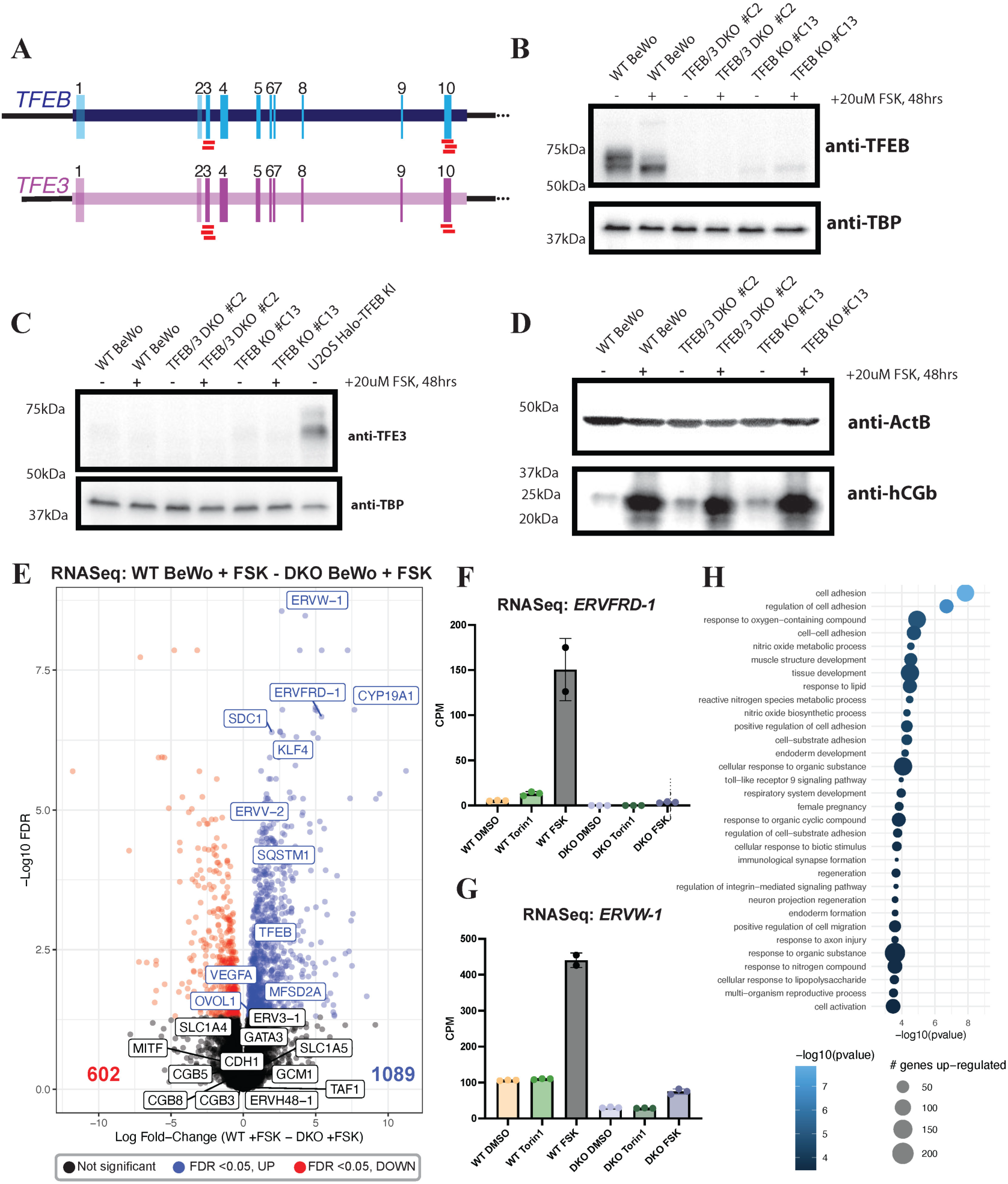
Perturbation of TFEB and TFE3 in BeWo cells using CRISPR KO. **A)** Strategy of CRISPR-based KO using sgRNAs (red segments) targeting the N-terminal and C-terminal regions of TFEB and TFE3 to induce gene deletion. **B)** Western blotting of TFEB in wild-type and CRISPR KO BeWo cells. **C)** Western blotting of TFE3 in wild-type, CRISPR KO BeWo cells and in U2OS with TFEB endogenously Halo-tagged as a positive control. **D)** Western blotting of syncytiotrophoblast marker hCGb in wild-type and CRISPR KO BeWo cells. **E)** Volcano plot of RNA-Seq data comparing Forskolin-treated wild-type BeWo cells with Forskolin-treated TFEB/TFE3 DKO BeWo cells. Genes that are significantly higher in the Forskolin-treated DKO cells are shown in red, and genes that are significantly lower in the Forskolin-treated DKO cells are shown in blue. **F)** Normalized CPM values from the RNASeq data showing expression of *ERVFRD-1* (Syncytin-2) **(F)** or *ERVW-1* **(G)** across samples. Data points indicate separate RNASeq replicates and error bars show standard deviation. **H)** GO Analysis of genes upregulated in Forskolin-treated WT BeWo compared to Forskolin-treated DKO BeWos. Top Biological Processes GO Terms are shown.

To identify genes that require TFEB for expression during BeWo differentiation, we performed RNA-Seq in wild-type and DKO #C2 BeWo cells treated for 48 hours with 0.1% DMSO or 20µM Forskolin to induce differentiation. To identify genes which may be responsive to nuclear TFEB, we also performed RNA-Seq in wild-type and DKO #C2 BeWo cells treated with 250nM Torin1 for 2 hours, an mTOR inhibitor shown to induce nuclear TFEB localization.^36,37^ Principal component analysis of three replicates for each condition showed that DMSO-treated and Torin1-treated samples clearly segregated from Forskolin-treated samples (Fig. S2A). Interestingly, differential gene expression analysis identified more than 1000 genes whose expression was significantly higher in the Forskolin-treated wild-type cells compared to the Forskolin-treated DKO BeWo cells (Fig. 1E). Among those genes most significantly dysregulated were known markers of syncytiotrophoblast differentiation including the syncytin loci *ERVFRD-1* (Syncytin-2) and *ERVW-1* (Syncytin-1) as well as syncytiotrophoblast-expressed genes *ERVV-2, SDC-1, KFL4,* and *CYP19A1* (aromatase) (Fig. 2E-G).^17,45^ RT-qPCR confirmed the severe defect in up-regulating *ERVFRD-1* in both the TFEB/3 DKO as well as the TFEB KO #c13 cells (Fig. S2B). Notably, GCM1 transcripts were not significantly affected in the RNA-Seq, suggesting that TFEB’s role is not directly upstream of GCM1 mRNA expression (Fig. 1E). Transcription of the hCG hormone cluster genes expressed in BeWo cells (*CB3, CGB5,* and *CB8)* were also not affected by TFEB loss, congruent with the unchanged protein expression measured by western blotting (Fig. 1D,E). GO classification of these ∼1000 genes were significantly enriched in categories relevant to cell-cell fusion including cell adhesion, cell-substrate adhesion, integrin signaling as well as female pregnancy and cellular compartment analysis indicated enrichment in extracellular matrix and cell membrane processes (Fig. 1H, S2C). While RNA-Seq, which measures steady-state transcription, identified only few deregulated genes after 2-hour treatment with Torin1, astoundingly, *ERVFRD-1* was one of the 30 up-regulated genes in Torin1-treated wild-type cells compared to control (Fig. S2D) suggesting a link between nuclear TFEB and syncytin expression in BeWo cells.

**Figure 2.**
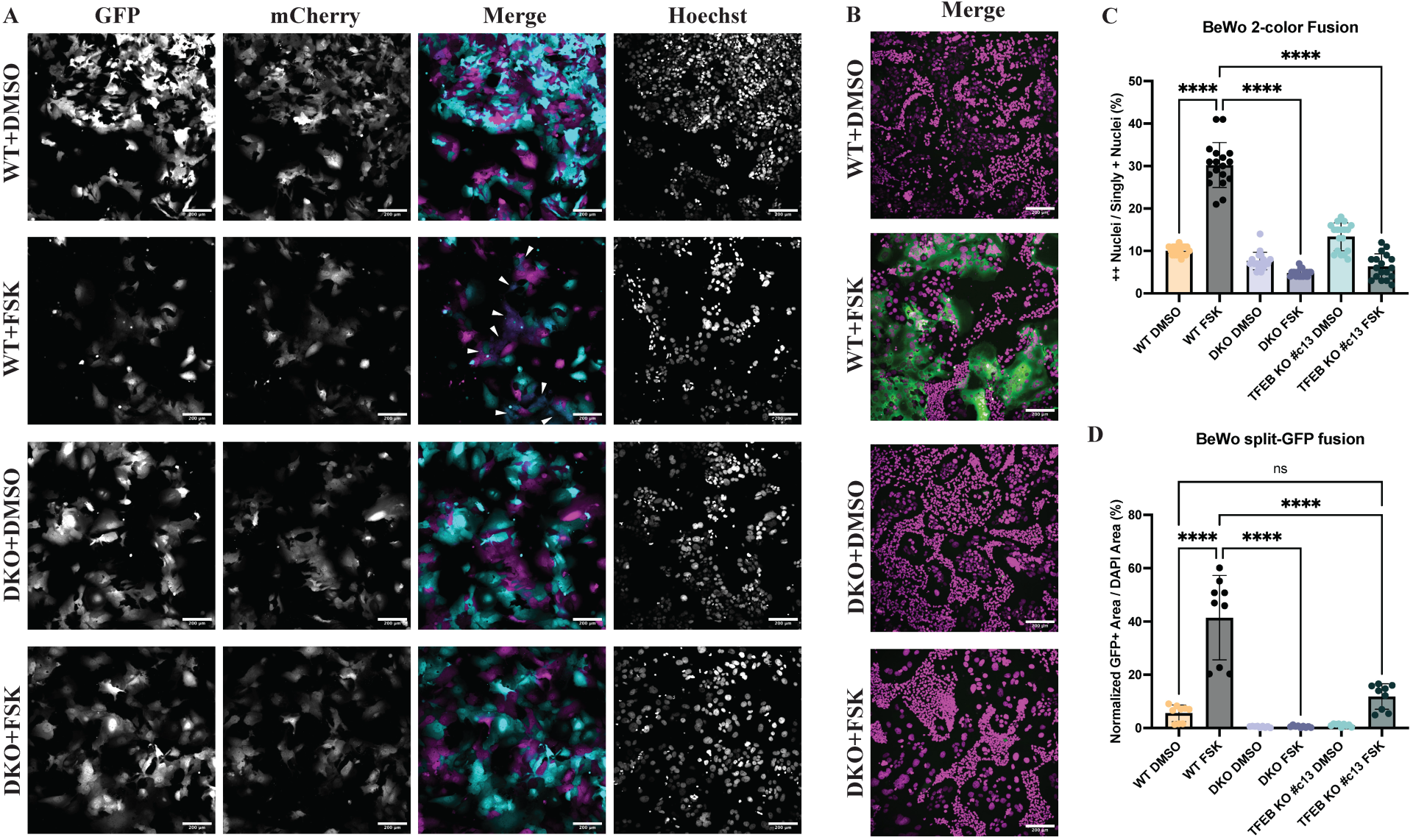
KO of TFEB/TFE3 in BeWo cells causes a functional defect in cell-cell fusion. **A)** Two color cell-cell fusion experiments co-culturing two populations of mCherry- and GFP-expressing BeWo cells were imaged on a spinning disk confocal microscope. Individual channels are shown in grayscale. In the merged composite image, the GFP channel is represented in cyan and the mCherry channel is represented in magenta. White arrows indicate fused syncytial areas in the Forskolin-treated wild-type cells. Scale bar = 200μm. **B)** Split-GFP cell-cell fusion experiments co-culturing GFP1-10 expressing BeWo cells with GFP11-expressing 293T cells and acquiring images on a spinning disk confocal microscope. In the merged composite image, the GFP channel is represented in green and the Hoechst channel is represented in magenta. Fused syncytial areas are shown by the reconstitution of GFP fluorescence shown in green. Scale bar = 200μm. **C)** Quantification of the two color cell-cell fusion experiments shown in A. **D)** Quantification of the split-GFP cell-cell fusion experiments shown in B. For C and D, statistical significance from an ordinary one-way ANOVA with Tukey’s multiple comparisons test is shown where ns = not significant,* = p<0.05, ** = p<0.01, *** = p<0.001, and **** = p<0.0001.

Given the apparent role for TFEB in syncytin expression, we next sought to determine whether the loss in syncytin transcripts results in a functional defect in cell-cell fusion. Two quantitative imaging-based assays were used to assess cell-cell fusion of wild-type, DKO, and TFEB KO BeWo cells upon 48-hour Forskolin-induced fusion. First, a two-color BeWo:BeWo fusion assay was performed by co-culturing BeWo cell lines Lentivirally transduced with either mCherry or GFP (Fig. 2A, S3A). Upon imaging of Hoechst-labeled nuclei in these two-color fusion experiments, quantification of the mCh/GFP double-positive nuclei over mCherry or GFP single-positive nuclei indicated the fused population (Fig. S3C). Forskolin treatment significantly increased the proportion of double-positive over singly-positive nuclei in the wild-type cells while this effect was nullified in the DKO and TFEB KO cells (Fig. 2C). In our RNA-Seq datasets, expression of the Syncytin-2 receptor MFSD2A was also lower in Forskolin-treated TFEB/3 DKO cells compared to wild-type, while the Syncytin-1 receptors SLC1A4/5 were unaffected (Fig. 1E). Because syncytins are unidirectional fusogens (i.e. they only need to be present on one membrane to induce fusion), loss of cell-cell fusion in a BeWo:BeWo fusion assay could be due either to loss of the fusogens themselves, Syncytin-1 or Syncytin-2, on one membrane or to loss of their receptors, SLC1A5 or MFSD2A, on the opposing membrane, respectively.^14^ To assess whether the TFEB-associated loss in syncytin expression alone significantly impacted cell-cell fusion, we also assessed cell-cell fusion between the wild-type, DKO, and TFEB KO BeWo cells with 293T cells. As observed by Mi et al., a donor cell line that constitutively expresses the necessary receptor proteins at high level (in their case MCF7, in our case 293T) can be seen to fuse and flatten into BeWo cells in a fully syncytin-dependent manner.^14^ In our experiment, we used a split-GFP approach where the respective BeWo cell lines were transduced with a Lentivirus to express GFP1-10 and co-cultured with 293T cells expressing GFP11. In wild-type cells, upon treatment with Forskolin, we observed large sheets of fused GFP-positive regions indicating cell-cell fusion between the 293T and BeWo cells, while in Forskolin-treated DKO and TFEB KO cells the GFP-positive area was dramatically reduced (Fig. 2B,D, S3B). Notably, there was a severely reduced capability of the TFEB KO #c13 cells to fuse with the 293Ts (Fig. 2D, S3B). We did observe a very small residual capacity of the TFEB KO #c13 cells to fuse with the 293Ts and attribute this to the small amount of remaining TFEB protein present in these cells (11%) that led to a small, but detectable increase in Syncytin-2 transcript levels upon Forskolin treatment (Fig. S2B). In sum, TFEB loss results in a severe defect in the normal differentiation-driven upregulation of syncytins that results in significant defects in cell-cell fusion.

TFEB’s main transcriptional role, as previously characterized in HeLa cells, is to control expression of many lysosomal and autophagy genes enriched in 10bp expanded E-box (CLEAR) motifs, termed the CLEAR gene network.^33,34^ Expression analysis of 11 CLEAR network TFEB target genes identified by Sardiello, *et al.* in our BeWo RNA-Seq dataset yielded only three which were meaningfully different between the Forskolin-treated wild-type and DKO cells, while most remained unchanged (Fig. 3A). Even though only three of the lysosomal genes were perturbed, in addition to gene expression data we next sought to investigate whether TFEB loss had any effect phenotypically on lysosomal organelle biogenesis. We stained BeWo cells with the live-cell pH-sensitive lysosomal marker Lysoview-540 (Biotium) and imaged live lysosomes in DMSO and Forskolin-treated BeWo cells (Fig. 3B). The number of lysosomes and the total area occupied by lysosomes were similar in the wild-type compared to the DKO and TFEB KO BeWo cells (Fig. 3C-D, S3D). Perhaps not surprisingly given our findings of TFEB in BeWo cells, lysosomes increased during Forskolin treatment regardless of TFEB loss with even larger gains in lysosomes in the TFEB DKO and TFEB KO #c13 cells than in wild-type cells (Fig. 3C-D). Thus, lysosomal biogenesis in BeWo cells is activated upon Forskolin differentiation but seems to be largely independent from TFEB expression, suggesting that an alternative or redundant factor may instead be controlling lysosomal gene expression in placental cells.

**Figure 3.**
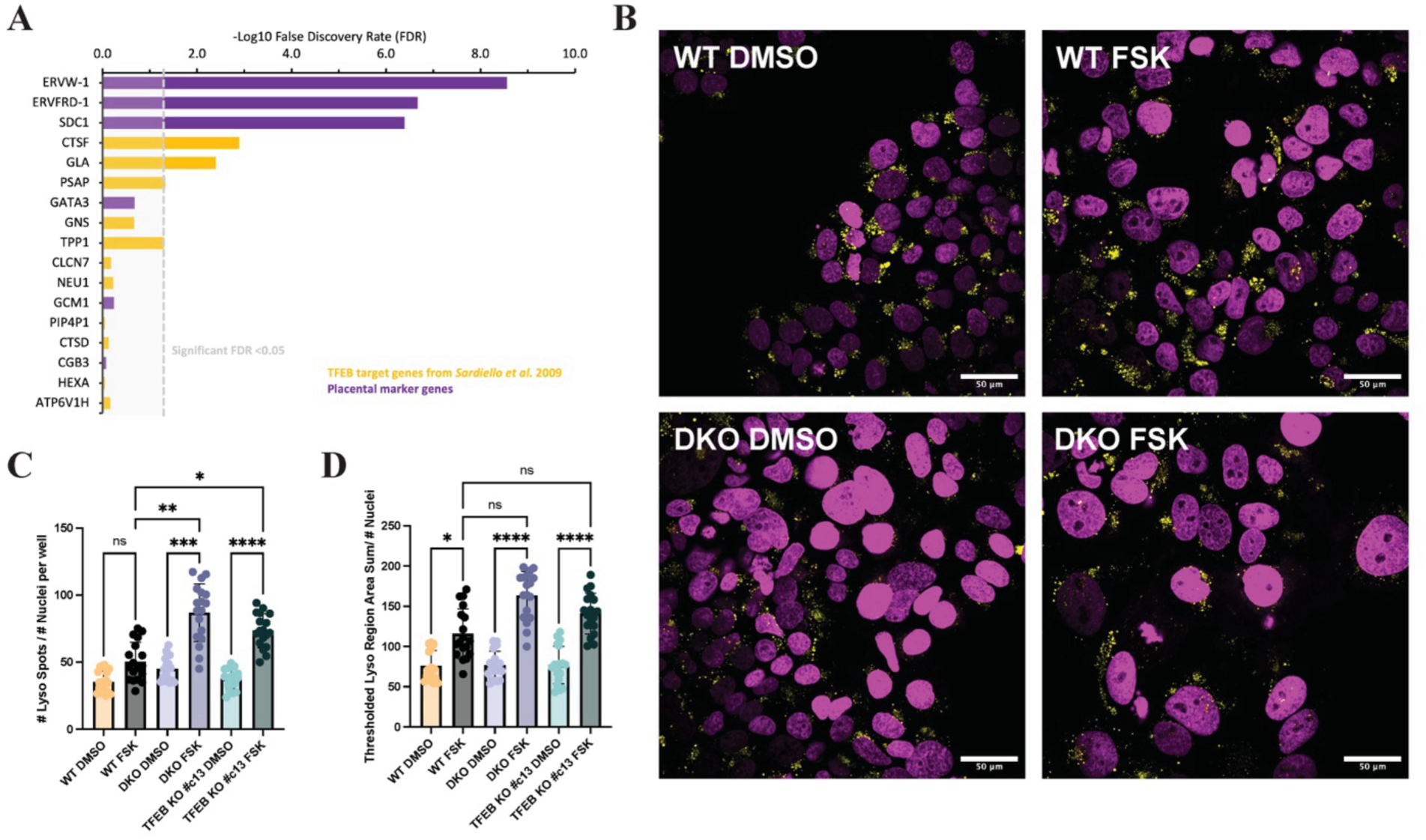
TFEB/3 KO in BeWo cells has minimal effects on lysosomal biogenesis or lysosomal gene expression. **A)** Differential gene expression analysis of previously identified lysosomal TFEB target genes (yellow bars) and selected syncytiotrophoblast marker genes (purple bars). The -Log10 False Discovery Rate of the comparison between Forskolin-treated wild-type BeWo cells and Forskolin-treated TFEB/3 DKO BeWo cells is shown with a significance cutoff of FDR = 0.05, equivalent to -Log10(FDR) = 1.30. **B)** Live lysosomes were imaged in DMSO-treated or Forskolin-treated BeWo cells by staining with Lysoview-540 and imaging on a spinning disk confocal. In the merged composite image, the Hoechst channel is represented in magenta and the Lysoview staining shown in yellow. Scale bar = 50μm. **C)** The normalized number of lysosomes was quantified by the per-well mean of total lysosomal spots detected per image divided by the total number of nuclei per image. **D)** The normalized total lysosomal area is quantified by the per-well mean of 540nm-positive thresholded region summed area per image divided by the total number of nuclei per image. For C and D, statistical significance from Kruskal-Wallis ANOVA test with Dunn’s multiple comparison test is shown where ns = not significant, p<0.05 = *, p<0.01 = **, p<0.001 = ***, and p<0.0001 = ****.

To more closely examine TFEB’s mechanism in controlling syncytiotrophoblast gene expression, we investigated TFEB’s nuclear and cytoplasmic partitioning during Forskolin treatment. We performed high-throughput confocal imaging of JF646-stained BeWo cells overexpressing Halo-3xFlag-TFEB driven by an L30 promoter and treated with DMSO or Forskolin. In untreated cells, TFEB localization is variable with most cells exhibiting cytoplasmic TFEB and some showing predominantly nuclear TFEB (Fig. 4A). Segmentation and quantification of the nuclear/cytoplasmic ratio of TFEB demonstrated a ∼2-fold increase (from ∼1 to ∼2) in nuclear/cytoplasmic TFEB after 48-hour treatment with Forskolin, similar to the effect of acute 1-hour Torin1 treatment (Fig. 4B). Western blot staining of TFEB in BeWo cells also showed a downward shift in the apparent molecular weight of the anti-TFEB band upon Forskolin treatment (Fig. 1B), a common signature of TFEB being dephosphorylated, an observation consistent with current evidence that TFEB’s shuttling to the nucleus is concomitant with its dephosphorylation.^36,37,46^

**Figure 4.**
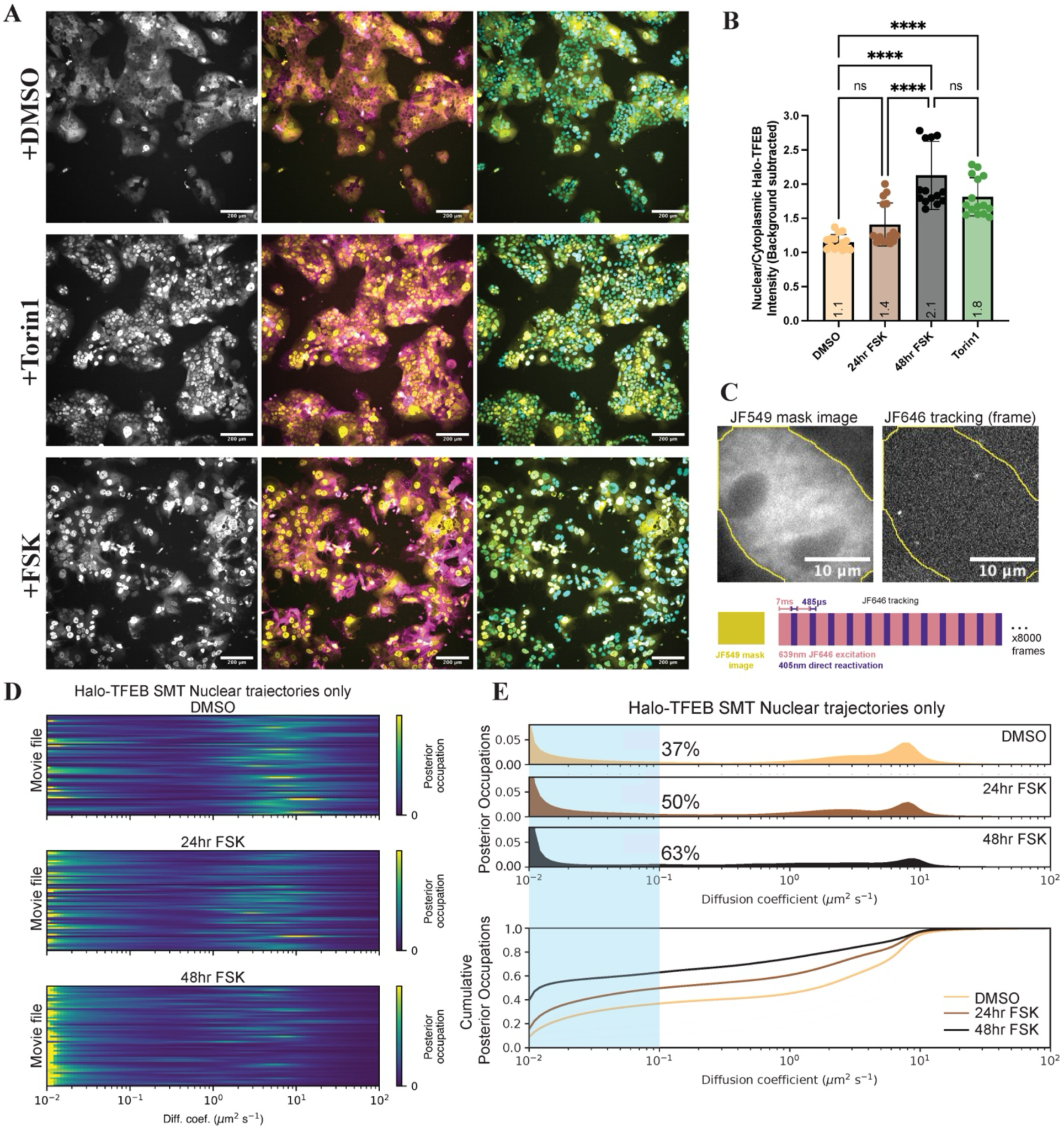
TFEB translocates into the nucleus and binds chromatin during syncytiotrophoblast differentiation. **A)** Live cell spinning disk confocal imaging of Halo-TFEB after treatment with Torin1 for 2hrs or treatment with Forskolin for 48hrs. The Halo-TFEB channel alone is shown in grayscale (left), followed by a composite image showing Halo-TFEB (yellow) with the membrane stain Biotium MembraneSteady-488nm (magenta) (middle), and Halo-TFEB (yellow) with Hoechst-stained nuclei (cyan). Scale bar = 200μm. **B)** Quantification of the nuclear/cytoplasmic ratio of Halo-TFEB in A. Statistical significance from an ordinary one-way ANOVA with Tukey’s multiple comparisons test is shown where ns = not significant, * = p<0.05, ** = p<0.01, *** = p<0.001, and **** = p<0.0001. **C)** Diagram of single molecule tracking of Halo-TFEB performed in BeWo cells. An example JF549 mask image (left) and a single frame from a JF646 tracking movie (right) are shown with the binarized mask outline used for segmenting nuclear versus external trajectories shown overlaid in yellow. Scale bar = 10μm. The bottom diagram shows how image acquisition was performed by acquiring a single JF549 image used for masking followed by stroboscopic illumination and activation of dark JF646 molecules using 405nm activation in the camera readout time. **D)** Heatmaps of diffusion coefficients following Bayesian analysis of single molecule trajectory data of Halo-TFEB in BeWo cells in different drug conditions. Each row corresponds to the distribution of posterior occupations for a single movie file. **E)** The distribution of diffusion coefficient occupancies for nuclear segmented trajectories of Halo-TFEB in different drug conditions. The fraction bound (calculated by the fraction of the distribution with a diffusion coefficient <0.1μm^2^/s) is shown highlighted with the blue region and the quantified values displayed as black percentages. The cumulative distribution function (CDF) of this same distribution is shown below.

Given TFEB’s redistribution to the nucleus upon Forskolin-treatment, we investigated whether TFEB’s chromatin binding within the nucleus was altered during differentiation. To sensitively measure TFEB-chromatin binding in live cells, we performed single molecule tracking (SMT) of BeWo cells stably overexpressing Halo-TFEB. SMT allowed us to assess the proportion of TFEB molecules engaged in chromatin binding (“bound”) versus freely diffusing (“free”) by collecting trajectories from ∼60,000 individual TFEB molecules across ∼60 cells per condition. We labeled Halo-TFEB overexpressing BeWo cells with two JF dyes in tandem: sparse (25nM) JFX646 for tracking and more dense (50nM) JF549 to assess the overall distribution of TFEB in the nucleus and cytoplasm. We then performed stroboscopic fast tracking (7ms frame rate) with interleaved direct photoreactivation of dark molecules using 405nm light to accurately capture fast moving molecules (Fig. 4C). Single molecule tracking of control constructs Halo-H2B (a highly chromatin-bound protein) and Halo-NLS (a diffusing control) yielded bound fractions of ∼85% and ∼9%, respectively, indicating the dynamic range of such an assay to detect immobile versus freely diffusing biological molecules (Fig. S4A,B, Supp. Movie 1, 2). Additionally, Forskolin-treatment alone did not induce a major change in the diffusion of the Halo-NLS or Halo-H2B controls (Fig. S4A,B). In DMSO-treated cells, Halo-TFEB diffusion modeled using saSPT exhibited a diffusion spectrum with approximately three modes: ∼19% of the population was immobile with a diffusion coefficient of ∼0.01μm^2^/s while the remaining molecules were diffusing with modes at diffusion coefficients of ∼2.5μm^2^/s and ∼9μm^2^/s (Fig. S4C,D, Supp. Movie 3). In cells treated with Forskolin for 24 hours, the fraction of immobile Halo-TFEB molecules increased from 19% to 42% and further raised to 62% in cells treated with Forskolin for 48hrs (Fig. S4C,D, Supp. Movie 4, 5). Since our acquired tracking movies included Halo-TFEB trajectories from both the nucleus and cytoplasm, two potential explanations arise from the observed increase in TFEB binding during Forskolin treatment. First, a simple explanation may be that Forskolin-induced relocalization of TFEB causes an increase in the proportion of nuclear trajectories thus increasing the total bound fraction, while the underlying nuclear behavior of TFEB remains the same. It could also be that during Forskolin induced differentiation, the nuclear population of TFEB also changes its behavior (via post-translational modifications, expression of cofactors, change in the underlying chromatin, etc.) which results in increased binding. To test these two possibilities, we used the collected JF549 images to create conservative, eroded masks of the nucleus in each movie and analyzed only trajectories which were entirely contained within the nuclear mask (Fig. 4C). Interestingly, in the nuclear-masked movies, the nuclear fraction of TFEB also significantly increased its bound fraction in Forskolin-treated relative to DMSO-treated cells, from 37% to 63%, respectively (Fig. 4D,E). Because the DMSO-treated cells contain many fewer trajectories within the nuclear masks than the Forskolin-treated cells, as a control we also re-ran the saSPT analysis after randomly sampling the same number of trajectories from each dataset, limited by the DMSO dataset that contained the fewest (∼20,000 trajectories) and saw no effect: we found agreement of the bound fraction within 1% indicating that our statistics are sufficiently powered (Fig. S5A-B). These data suggest that Forskolin differentiation induces not only a redistribution of TFEB from the cytosol to the nucleus but also an increase in its propensity to associate with chromatin once inside the nucleus.

To determine whether TFEB’s increased chromatin association is Forskolin-dependent, we tested TFEB nuclear localization and binding by SMT under other pharmacological perturbations. Consistent with previous reports, we measured increased TFEB nuclear localization upon treatment with either Torin1 or sucrose and Leptomycin B (Fig. S6A).^34,47^ In both cases we also measured an increased association with chromatin of the masked nuclear Halo-TFEB population, indicating that TFEB’s increased propensity for chromatin association is not necessarily Forskolin-dependent, but that increased chromatin binding follows from TFEB’s relocalization to the nucleus, regardless of chemical stimuli (Fig. S6B,C, Supp. Movie 6, 7).

Finally, we sought to measure which genomic sites this increasingly chromatin-associated TFEB may be binding to directly during differentiation. We performed ChIP-Seq of DMSO-treated, 2-hour Torin1-treated, and 48-hour Forskolin-treated stably expressing 3xFLAG-Halo-TFEB BeWo cells using a mouse anti-FLAG antibody. In agreement with a strong role of TFEB in stimulating the expression of the two syncytins governing cell-cell fusion, ChIP data shows clear TFEB enrichment at the promoter of both *ERVFRD-1* and *ERVW-1* upon Forskolin and Torin1 treatments (Fig. 5A-B). For subsequent analysis, two biological replicates were merged and statistically robust peaks identified with IDR analysis (Fig. S7A-C, E).^48^ Of these robust peaks, significant overlap exists between peaks identified in the Forskolin and Torin-1-treated cells with a larger number of peaks identified as Torin-1-specific (Fig. 5C). A de-novo MEME-ChIP motif search for TFEB peaks across all conditions confirmed the canonical TFEB motif, an expanded and somewhat flexible E-box motif, centrally enriched around TFEB peak summits (Fig. 5D-E).^49^ Differential TFEB binding after Forskolin and Torin1 treatment was also analyzed using MACS2 bdgdiff which identified three categories of peaks (Fig. 5F).^50^ Differential Forskolin-enriched peaks were few (530); most TFEB peaks were Torin1-enriched (6,312) or Common to both Torin1and Forskolin (5,343). These differentially enriched peaks were intersected with IDR robust peaks resulting in 2,176 Torin1-enriched peaks and 2,242 Forskolin and Torin1 Common IDR-robust peaks which were mapped to their nearest genomic element for analysis (Fig. S7D). Interestingly, some differences emerged in the GO ontology between the mapped genes for Torin1-enriched peaks and the Common peaks. Notably, both pathways seemed to converge on GTPase regulation, while Torin1-enriched peaks were also enriched in metabolic processes such as endocytosis and vesicle organization and Common peaks pointed to cell adhesion, regulation by Wnt signaling, and autophagy (Fig. 5G). Interestingly, promoter-proximal TFEB peaks were enriched for the more canonical lysosomal and autophagy regulatory GO categories, while more distal TFEB peaks were enriched in Wnt signaling, cell adhesion, cell junction assembly, and morphogenesis suggesting that TFEB’s distal regulation may be critical for its role in syncytiotrophoblast differentiation (Fig. S7G-H).

**Figure 5.**
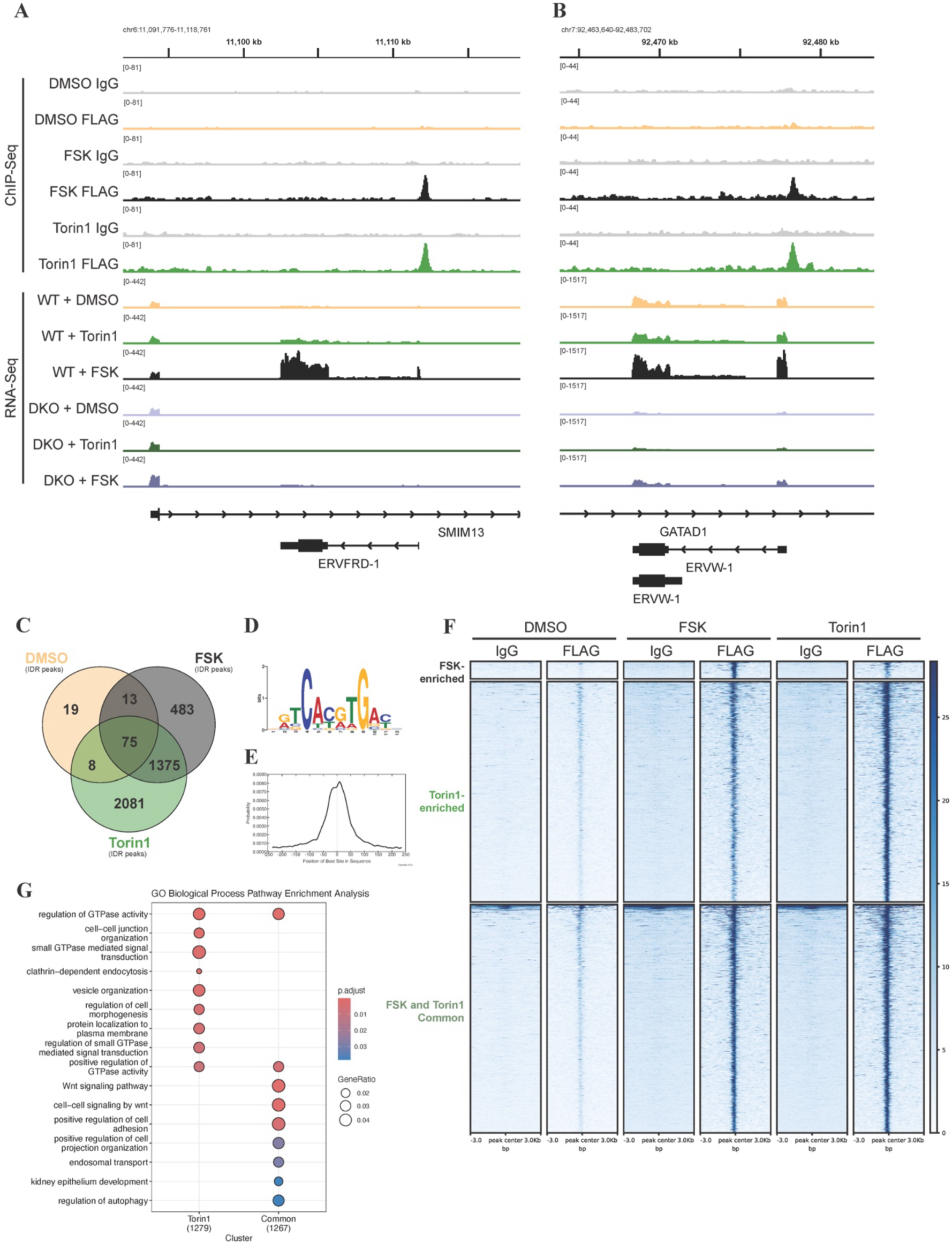
TFEB directly binds the *ERVFRD-1* and *ERVW-1* promoters to regulate their expression. Gene tracks displaying binned reads of 3xFlag-Halo-TFEB ChIP-Seq and RNA-Seq at the *ERVFRD-1* **(A)** and the *ERVW-1* **(B)** locus. Merged Replicates 1 and 2 are shown. **C)** Venn diagram of statistically significant 3xFlag-Halo-TFEB ChIP-Seq peaks, called with IDR Analysis (IDR score < 0.05) in each treatment condition. **D)** Top significant motif from MEME-ChIP analysis of significant 3xFlag-Halo-TFEB ChIP-Seq peak summits (IDR score <0.05) expanded by 250bp in both directions and pooled from all treatment conditions. **E)** Distribution analysis of the top MEME motif “DRTCACGTGAYH” in the analyzed sequences. **F)** Heatmaps of differentially enriched peaks analyzed with MACS2 bdgdiff showing +/-3000bp centered around each peak. Replicates 1 and 2 were merged and IDR robust peaks (IDR score <0.05) are shown, please see Fig. S7 for expanded plot of replicates. **G)** GO Enrichment Analysis of all annotated genes associated with at least one significant 3xFlag-Halo-TFEB ChIP-Seq peak (IDR score <0.05). Genes with higher Torin1 binding or with equivalent Forskolin and Torin1 (Common) enrichment (determined by MACS2 bdgdiff) are analyzed separately. Top Biological Processes GO Terms are shown.

Finally, since we have shown that 2-hour Torin1 treatment already leads to rapid nuclear accumulation of TFEB (Fig. 4B), increased Syncytin-2 transcript levels (Fig. S2D), increased TFEB-chromatin association by fast SMT (Fig. S6B), and binding of TFEB at relevant syncytiotrophoblast genes by ChIP (Fig. 5A-B), we hypothesized that the faster relocalization of TFEB in Torin1-treated over Forskolin-treated cells could result in faster rates of cell-cell fusion. Indeed, co-treatment with Torin1 and Forskolin resulted in a synergistic effect at 24-hours with a significantly higher proportion of fused cells than Torin1 or Forskolin treatment alone (Fig. 6A-B). We also found that Torin1 alone resulted in a significant increase in cell fusion, similar to that of Forskolin, at 48 hours (Fig. 6C-D). At 48 hours, Torin1 and Forskolin together increased fusion even more than either drug alone, however we noticed that with this high rate of cell fusion, cells began to detach from the dish. The data is consistent with a model that nuclear TFEB is sufficient for cell-cell fusion and that a Forskolin-dependent process synergizes with nuclear TFEB during early timepoints of cell-cell fusion.

**Figure 6.**
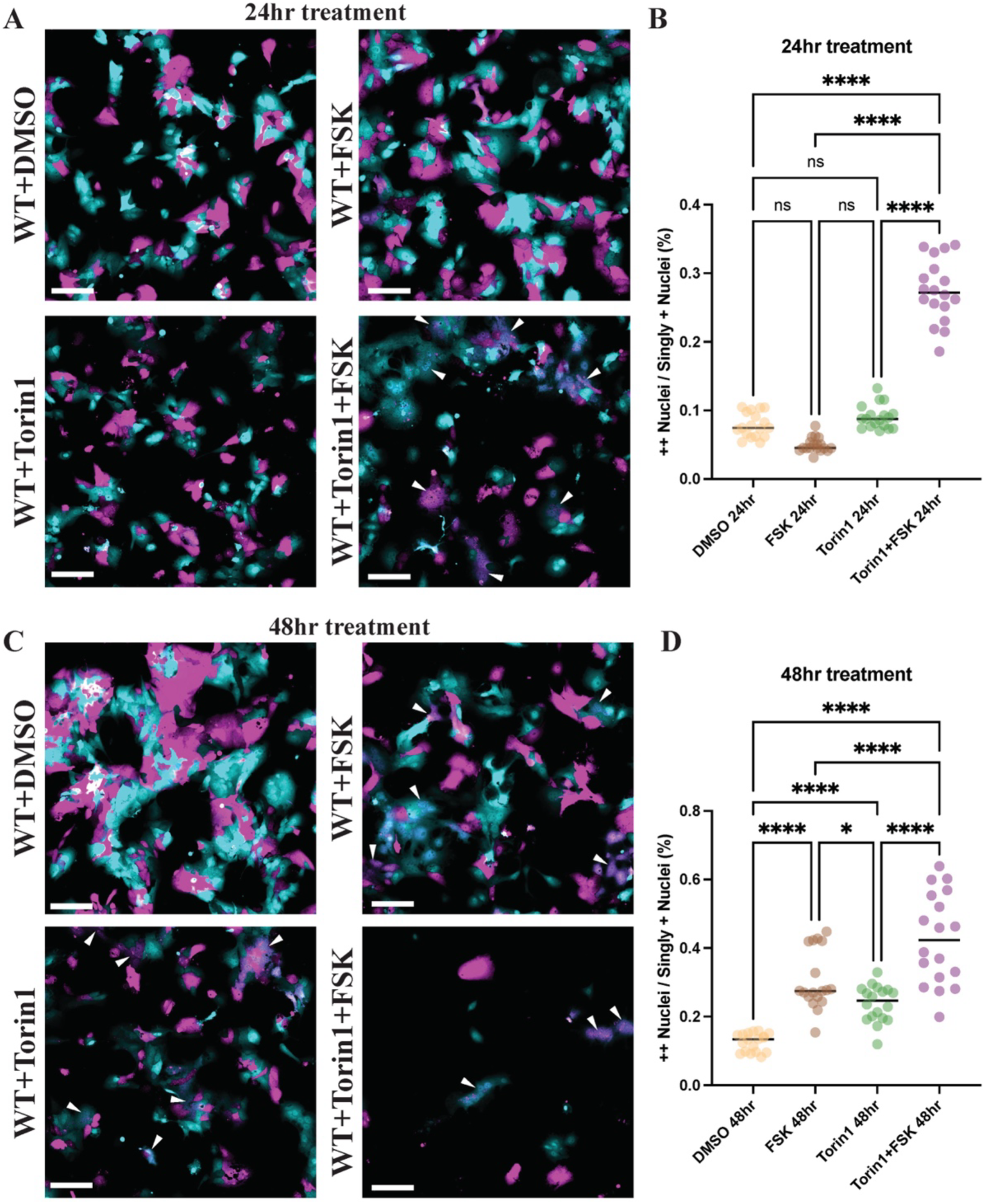
Torin1 treatment increases the rate of syncytiotrophoblast fusion. **A)** Two color cell-cell fusion experiments co-culturing two populations of mCherry-and GFP-expressing BeWo cells were treated with the indicated compounds for 24 hours, then imaged on a spinning disk confocal microscope. In the merged composite image, the GFP channel is represented in cyan and the mCherry channel is represented in magenta. White arrows indicate fused syncytial areas in the Torin1 and Forskolin-treated wild-type cells. Scale bar = 200μm. **B)** Quantification of the two color cell-cell fusion experiments shown in A. **C)** Same as A, except cells were treated with indicated compounds for 48hours. **D)** Quantification of the two color cell-cell fusion experiments shown in C. Statistical significance from an ordinary one-way ANOVA with Tukey’s multiple comparisons test is shown where ns = not significant,* = p<0.05, ** = p<0.01, *** = p<0.001, and **** = p<0.0001.

## Discussion

In our present study, we have identified a new direct regulator of *ERVFRD-1* and *ERVW-1* expression, genes encoding Syncytin-1 and -2 that play an essential role in cell-cell fusion of syncytiotrophoblasts. We demonstrated that TFEB is the predominant TFE family member expressed in BeWo cells and genetic loss of TFEB creates a significant loss in Syncytin-1 and -2, as well as MFSD2A expression upon differentiation, which leads to a loss in the ability for these cells to fuse (Fig. 1, 2). We further demonstrate in heterologous fusion assays that the Syncytin loss itself is sufficient for the loss in fusion in TFEB KO cells (Fig. 2). In producing TFEB KO clones, our highest knockout efficiency clone was still not 100% complete and showed some residual upregulation of *ERVFRD-1* upon Forskolin treatment (Fig. 1B, 1F). We postulate that this residual activity in comparison to the complete TFEB/TFE3 DKO clone is due to the small, yet detectable residual level of TFEB present, rather than the presence of TFE3, which was very lowly expressed at the mRNA level (RPKM <5) and undetectable by western blotting (Fig. 1). Lending strength to this argument are multiple lines of evidence that point to TFEB, rather than TFE3, playing an important role in placental biology. While homozygous TFEB KO in mice is lethal due to placental phenotypes, homozygous TFE3 or MITF KOs show no apparent gestational phenotype.^51^ A very recent study published during preparation of this manuscript further identified TFEB, and not TFE3, as a genetic hit in a CRISPR screen of genes required for syncytiotrophoblast formation in hTSCs.^52^ Finally, in single cell sequencing of human morula-stage embryos, TFEB was identified as a TE-enriched gene which correlated with the expression of GATA3, a trophectoderm marker, while TFE3 did not.^53^

Surprisingly, our RNA-Seq data identified dysregulated genes that point to a new role for TFEB in controlling expression of key syncytiotrophoblast genes, rather than its canonical lysosomal and autophagic gene targets (Fig. 1). Given the seeming importance of lysosomal and autophagy biology in placental cell differentiation, we were surprised by this finding.^54,55^ In our study, even double KO of TFEB and TFE3 did not perturb many lysosomal and autophagy genes or the ability to produce lysosomes during differentiation (Fig. 3). However, ChIP-Seq did identify TFEB binding with promoter-proximal enrichment at many lysosomal and autophagic target genes, as previously characterized, as well as binding to many key syncytiotrophoblast genes. There thus seems to be a clear disconnect between TFEB’s genetic targets and its transcriptional functions. Interestingly, several other recent publications lend credence to the idea of cell-type dependent roles of TFEB outside of its role in CLEAR network regulation. In undifferentiated mouse ESCs, TFEB instead binds to the *Nanog* promoter and controls pluripotency genes, while TFE3 rather than TFEB controls the canonical lysosomal and autophagy network genes.^56^ In zebrafish, *tfeb/tfe3a/tfe3b* triple mutant animals showed almost no changes to steady state levels of autophagy or lysosomal genes; TFEB-dependence of these genes was only observed upon immunological stress.^57^ Most interestingly, Wnt-driven nuclear localization of TFEB results in an activated subset of Wnt-driven TFEB genes, including genes involved in cell adhesion and secretion, being activated without perturbing expression of lysosomal target genes or lysosomal biogenesis.^58^ These results are consistent with our finding that Torin1 and Forskolin-enriched common genes were enriched in GO categories for Wnt signaling and modulation of cell adhesion (Fig 5G). Overall, these results lead to several interesting hypotheses for TFEB’s altered role in placental cells: it may be that a key TFEB cofactor for regulating the CLEAR network is not present in placental cells and thus TFEB can bind, but not transactivate, these classical targets. It could also be that placental-specific post-translational modifications alter TFEB’s transactivation abilities at specific genes (as is the case for PARsylated Wnt-driven TFEB targets).^58^ It is possible that another E-box binding factor (outside of the TFE family) is present in trophoblast cells and instead controls the canonical CLEAR network through direct competition with TFEB at these sites. These possibilities will be interesting to distinguish in future studies.

Biophysically, we have shown that nuclear import of TFEB leads to increased chromatin engagement regardless of the chemical initiator of TFEB import (i.e. Forskolin, Torin1, or Sucrose and Leptomycin B) (Fig. 4). This quantitative finding suggests that as nuclear TFEB concentration increases, the proportion of TFEB bound to chromatin also increases. Due to the limited number of trajectories collected per cell (n∼200) a cell-by-cell analysis of chromatin-binding versus TFEB concentration was not possible. However, we speculate that because TFEB binds DNA as a homo- or hetero-dimer that this increased propensity to bind DNA as a function of concentration may simply be a reflection of the bias towards dimerization at higher nuclear protein concentrations.^31^ In addition, we have shown that Torin1 treatment synergizes with Forskolin to increase the rate of cell-cell fusion (Fig. 6). These data suggest that the concentration of nuclear TFEB may be an important rate-limiting step in the induction of cell-cell fusion. While nuclear TFEB localization with Torin1 treatment alone was alone sufficient to induce cell-cell fusion, given Torin1’s inability to increase cell-cell fusion by 24hours, it is likely that nuclear TFEB is upstream from a second, rate-limiting process which is Forskolin-dependent.

In our study we have identified TFEB as a novel regulator of Syncytin-2 expression, and one which appears orthogonal to existing regulation by GCM1. We found that TFEB depletion did not affect the expression of GCM1 at the transcript level and both GCM1 and TFEB bind distinct DNA motifs to regulate syncytin.^23,59^ Yet, ChIP-Seq of hTSCs differentiated to STB shows GCM1 occupancy at ∼300bp upstream of the *ERVFRD-1* promoter, which overlaps with the TFEB peak identified here in our ChIP-Seq.^52^ Genetically, both individual GCM1 and TFEB KO cells individually are defective in cell-cell fusion, where presumably the other factor is expressed, thus suggesting that both are required for syncytiotrophoblast fusion. It is not yet known whether TFEB may physically interact with GCM1. TFEB protein-protein interaction studies have not yet been performed in trophoblast cells and since GCM1 is almost exclusively expressed in the placenta^60^, any putative interaction has not been documented but will be interesting to investigate.

Finally, therapeutic modalities to prevent or treat pregnancy diseases are desperately needed and our study may provide a unique therapeutic angle on treating diseases where cell-cell fusion is perturbed, such as preeclampsia and fetal growth restriction. Research is divided on how lysosomal and autophagy processes may contribute to placental defects in pregnancy disorders. Elevated LC3 levels and decreased p62/SQSTM1 levels were found in western blotting of placentas from patients diagnosed with severe preeclampsia.^61–63^ On the other hand, Nakashima *et al.* found elevated levels of p62/SQSTM1 in EVT cells from preeclamptic placentas, but not in the syncytial villous trophoblast suggesting that autophagy inhibition in a subset of cells, rather than activation more broadly, may contribute to disease.^64^ At the transcriptomic level, a comparison of five microarray datasets of primary placental tissues found that the gene expression of macro-autophagy genes (GO:0016236) was not significantly different between healthy and preeclamptic placentas. On the other hand, drugs that affect TFEB phosphorylation and localization such as Torin1, Rapamycin, and amino acid shortage have shown to have direct effects on syncytiotrophoblast formation. However, it was previously unclear which factor in the pathway was important for enabling the effects of these pleiotropic drugs.^10^ Upon identifying TFEB as a direct transcriptional effector in trophoblast fusion, this study may reconcile some of these findings by separating TFEB’s importance in syncytiotrophoblast differentiation from its role in controlling lysosomal and autophagy genes. Further investigation is warranted since our findings predict that controlling TFEB directly either via genetic perturbations or via targeted pharmaceutical interventions may offer a novel approach to regulating cell-cell fusion during syncytiotrophoblast development.

## Materials and Methods

### BeWo cell culture

Human placental choriocarcinoma BeWo cells (ATCC, CCL-98) were cultured at 37 °C and 5% CO_2_ in F12-K medium (Corning or ATCC) supplemented with 10% fetal bovine serum and 10 U ml^−1^ penicillin-streptomycin. Cells were passaged by trypsinization every 2-4 days and split at a ratio of 1:3-1:5. Media was changed completely every 24-36hrs.

#### Treatments

As indicated in the text, BeWo cells were treated with 20µM Forskolin dissolved in DMSO (Millipore Sigma F3917), 250nM Torin1 (Cell Signaling Technology #14379S), or 0.1% DMSO in supplemented F-12K medium. For sucrose treatment, crystalline D-Sucrose (Fisher #BP220-1) was dissolved in supplemented F-12K medium to 100mM final concentration, 0.2µm filtered, then added to cells in culture. As indicated, LeptomycinB (InvivoGen #inh-lep-10) dissolved in ethanol was used at 20nM final concentration in 100mM sucrose supplemented F-12K medium.

### Cas9 Editing

#### Plasmid method (TFEB/3 DKO #C2)

BeWo cells were plated at 300,000 per well into a 6-well plate and transfected with ten pU6 sgRNA CBh Cas9 PGK Venus plasmids carrying guides targeting TFEB and TFE3 (Table 1) with Lipofectamine 3000 per the manufacturer’s instructions. Two days post transfection, Cas9-Venus+ cells (∼6% of the total population) were FACS sorted into 96-well plates. Cells were grown for ∼2-3 weeks and then genotyped for genomic deletions in TFEB and TFE3 by PCR (Table 2). Clone #C2 was validated by western blotting and Sanger sequencing (Table 3).

**Table 1.**
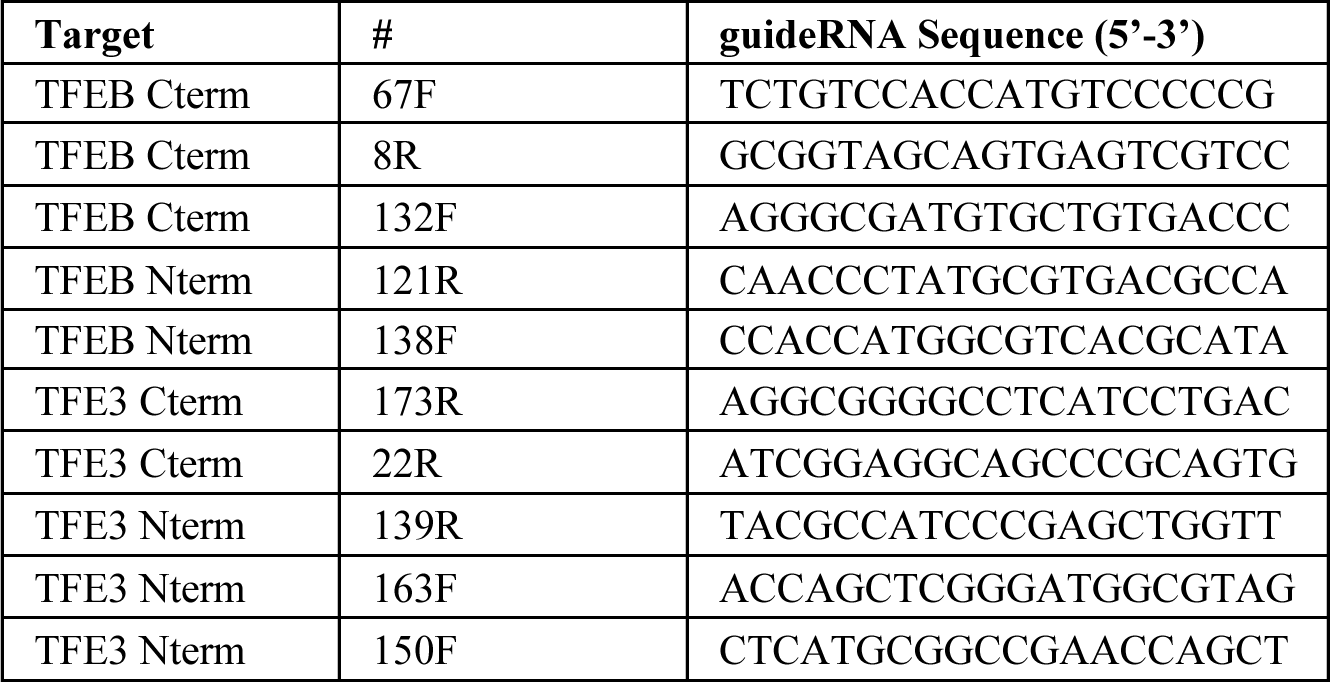
GuideRNAs for TFEB/TFE3 KO.

**Table 2.**
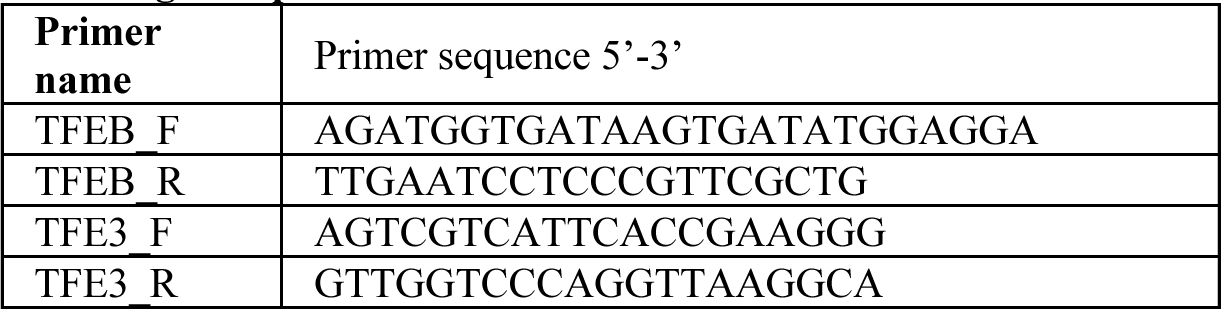
gPCR primers for TFEB/TFE3 DKO.

**Table 3.**
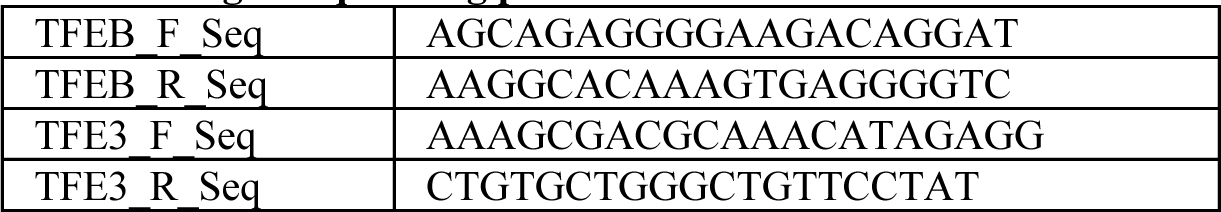
Sanger sequencing primers for TFEB/TFE3 DKO.

#### RNP Nucleofection (TFEB KO #c13)

Guide RNAs were ordered from IDT. Cas9 RNP was assembled on ice by adding 0.7µL160uM crRNA + 0.7µL160uM trRNA in a PCR strip tube and incubated in a thermocycler at 37C for 30min. Then, 1.4µL 40uM purified Cas9-NLS (QB3 Macrolab) was added to the duplexed sgRNA and incubated in a thermocycler at 37C for 15min. BeWo cells were split and counted using Trypan blue dead-cell exclusion, and 3E5 cells per reaction were aliquoted into a 15mL conical and centrifuged at 200g for 2.5min. Cells were resuspended in 20µL per reaction of Lonza SG supplemented solution per the manufacturer’s instructions. 2.5µL of complexed Cas9 RNP was added to 20µL BeWo cells in Lonza SG solution and transferred to a Lonza 16-well strip nucleofection cuvette then nucleofected with a Lonza 4D Nucleofector using program CA-137. Immediately following nucleofection, 75µL of warm F-12K media was gently added to the top of the BeWo cell reaction without mixing and the cells were allowed to recover for 10min at 37C, 5% CO2. After recovery, cells were then transferred from the cuvette into a 6-well plate for growth.

### Western blotting

Cells were trypsinized, split, and counted and 1E6 cells were centrifuged at 200g for 2.5min then resuspended in 150µL PBS + 50µL 4x SDS-PAGE loading buffer (200mM Tris pH 6.8, 400mM DTT, 10% B-ME, 8% SDS, 0.4% bromophenol blue, 40% glycerol) and boiled at 100°C on a heat block for 20 minutes and then centrifuged at 16,000g (Eppendorf 5415) for 3 minutes at 4°C. Supernatant was then transferred to a fresh tube and stored at -20°C. 10μL of cell lysate was loaded onto 4-20% Tris-glycine (BioRad) and transferred onto 0.45um nitrocellulose. A solution of 5% bovine serum albumin was used to block membranes for 1 hour at room temp prior to blotting and used to dilute primary and secondary antibodies. Gels were imaged by addition of Western lightning ECL reagent on a Chemidoc imaging system.

#### Antibodies

ACTB, used at 1:5000, Sigma Aldrich #A2228; TBP, used at 1:2000, Abcam #ab51841; hCGb, used at 1:1000, Abcam #ab53087; TFEB, used at 1:1000, Cell Signaling Technology #4240S; TFE3, used at 1:5000, Abcam #ab93808; Anti-rabbit HRP secondary, used at 1:5000, Invitrogen #31462; Anti-mouse HRP secondary, used at 1:5000, Invitrogen #31430.

### RNASeq

BeWo cells were plated at 300,000 cells per well into a 6-well plate and allowed to settle 6 hours to overnight. Cells were then treated with 0.1% DMSO or 20uM Forskolin for 48 hours. For Torin1 treatment, cells were treated with 0.1% DMSO for 42 hours and then switched to 250nM Torin1 media for 2 hours before harvesting. Media for all treatments was changed at 24 hours. Three biological replicates were performed by plating and treating cells three different times. RNA was extracted from each well using 500µL Trizol followed by two sequential chloroform extractions followed by ethanol precipitation per the Trizol User Guide (Thermo Fisher #MAN0001271). Total RNA was depleted of rRNA using the Illumina rRNA Depletion Kit (NEB #E6310) and then prepared for Illumina sequencing using the NEBNext Ultra II Directional RNA Library Prep Kit for Illumina #E7760 per the manufacturer’s protocol. Sequencing was performed by MedGenome on an Illumina NovaSeq with paired-end 150bp reads. Fastq files were assessed by FASTQC, then pseudo-aligned to the hg38 transcriptome using Salmon to quantify transcripts in mapping-based mode (salmon quant -i) using decoy-aware mapping with gencode.v39.transcripts.fa. Differential gene expression analysis was performed with EdgeR in Jupyter Notebook. Salmon-mapped transcripts were imported into Jupyter Notebook in R using tximport. Transcript counts were normalized for both sequencing depth and effectively normalized against highly variable transcripts using the edgeR function calcNormFactors. Transcripts were then filtered by expression across all samples using edgeR filterByExpr(min.count = 10, min.total.count = 15). Dispersion was estimated using estimateDisp with classic mode, then statistical comparisons between individual sample conditions were made using EdgeR makeContrasts then glmQLFit. The reported p-value and FDR values were corrected with adjust.method = “BH” and differentially expressed genes are reported with a adjusted FDR<0.05.

### qPCR

RNA was harvested from cells using Trizol followed by chloroform extraction and ethanol precipitation. 1μg RNA from each sample was then reverse transcribed using iScript (BioRad) following the manufacturer’s instructions. Amplification was performed using CFX master mix (Biorad) and a 2-step PCR cycling protocol with 60°C annealing. All qPCR primers were evaluated using a dilution series to ensure calculated primer efficiencies of 90-110% and amplicons were validated using agarose gel electrophoresis and Sanger sequencing.

### Cell-Cell Fusion Assays

#### Stable cell lines

To create stable cell lines, BeWo cells were grown to ∼50% confluency in a 6-well plate and transduced with 500µL crude Lentivirus (pHAGE EF1a mCherry IRES Hygro, pHAGE EF1a GFP IRES Hygro, or SFFV GFP1-10 IRES Hygro) in F12-K supplemented media with 0.8μg/mL polybrene. Lentivirus was incubated with the cells for 16-24hrs, then media was changed. Stable cell lines were created by selecting with 200μg/mL Hygromycin for 2 weeks and maintained in Hygromycin selection conditions. To create stable HEK293T cells, HEK 293T cells were plated to ∼75% confluence in a 6-well plate and transfected with 5μg pQCXIP CMV BSR-GFP11 IRES Puro (BSR = Blasticidin Resistance) with 7.15µL PEIMax in serum free DMEM. Two days post transfection, cells were selected in 8μg/mL Blasticidin for 2 weeks and maintained in antibiotic selection conditions. Plasmids used are listed in Table 4.

**Table 4.** Plasmids.

| Reagent | Experiment used | Source |
| --- | --- | --- |
| pHAGE L30prom H2B-GDGAGLIN-Halo-V5 IRES Neo | Single molecule tracking | Tjian-Darzacq lab |
| pHAGE L30prom Halo-NLS IRES Puro | Single molecule tracking | Tjian-Darzacq lab |
| PiggyBac L30 3xF-Halo-GDGAGLIN-TFEB IRES Puro | Single molecule tracking | Tjian-Darzacq lab, cloned from TFEB plasmid gifted by Zoncu lab |
| pHAGE EF1a mCherry IRES Hygro | Fusion assays | Tjian-Darzacq lab |
| pHAGE EF1a eGFP IRES Hygro | Fusion assays | Tjian-Darzacq lab |
| pHR-SFFV-GFP1-10 plasmid | Fusion assays | Addgene, #80409 |
| pQCXIP-BSR-GFP11 | Fusion assays | Addgene, #68716 |
| pU6 sgRNA CBh Cas9 PGK Venus | CRISPR editing | Tjian-Darzacq lab, derivative of Zhang lab plasmid |
| pMAX-GFP | Control | Lonza |

For 2-color fusion experiments, EF1a mCherry IRES Hygro and EF1a GFP IRES Hygro expressing BeWo cells were split and mixed at a 1:1 ratio so that 10,000 cells (5,000 mCherry + 5,000 GFP) were plated per well of a 96-well plate into Perkin Elmer Cell Carrier Ultra plates. For splitGFP fusion, 10,000 SFFV GFP1-10 IRES Hygro BeWo cells were plated into Perkin Elmer Cell Carrier Ultra plates. BeWo cells were allowed to settle and adhere for 3 hours to overnight, then media was aspirated and changed to 100μL/well of F12-K supplemented media containing 0.1% DMSO (control) or 20uM Forskolin (treatment). For splitGFP experiments, immediately after changing the media on the GFP1-10 BeWo cells to DMSO or Forskolin, 293T cells stably expressing pQCXIP CMV Blasticidin-GFP11 IRES Puro were split, centrifuged at 200g for 2.5min, and resuspended in a small volume of 293T media (4.5g/L DMEM containing Glutamax, Sodium Pyruvate, and PenStrep) and counted using Trypan blue dead-cell exclusion. 5,000 293T BSR-GFP11 cells in a volume of 1-5µLwere added on top of the BeWo cells. Cells were treated with 0.1% DMSO or 20uM Forskolin for a total of 48hrs before imaging and media was changed every 24hrs. Cells were then imaged on a spinning disk confocal microscope with 10X Air objective.

#### Confocal Imaging

Immediately before imaging, media was changed to 4.5g/L DMEM without phenol-red containing 10% FBS, 1x Glutamax supplement, and 1x Sodium Pyruvate supplement plus a final concentration of 2μg/mL Hoechst 33342 (Invitrogen) and incubated for 10-15min before beginning imaging. Plates were sealed with Sigma Breathe-Easy sealing membranes. High throughput spinning disk confocal imaging was performed on a Perkin Elmer Opera Phenix microscope with incubation at 37C with 5% CO2. Two Peak Autofocus and Confocal Mode imaging were used and no binning was performed during acquisition.

For BeWo 2-color fusion, a 10x Air NA=0.3 objective was used and the 4 channels were acquired (Brightfield, EGFP, mCherry, and Hoechst) where Hoechst/Brightfield were collected in the same exposure and separated from mCherry/GFP which were collected in a second exposure. The following illumination settings and filter sets were used: Brightfield Transmission, Emission 650-760nm, 100ms exposure at 20% power; GFP 488nm Excitation, 500-550 Emission, 1000ms exposure at 100% power; mCherry 561nm Excitation, 570-630nm Emission, 1500ms exposure at 100% power; Hoechst 375nm Excitation, 435-480nm Emission, 300ms exposure at 100% power. Z-stacks with 2-3 planes and 7um separation were taken to ensure in-focus images were taken across the plate and 9 central fields of view were acquired per well which covered almost the entire well area.

For splitGFP cell-cell fusion imaging, a 10x Air NA=0.3 objective was used and the following laser and filter sets were used to acquire three channels: Brightfield Transmission, Emission 650-760nm, 100ms exposure at 50% power; GFP 488nm Excitation, 500-550 Emission, 2000ms exposure at 100% power; Hoechst 375nm Excitation, 435-480nm Emission, 400ms exposure at 100% power. Hoechst/Brightfield were collected in the same exposure and separated from GFP which was collected in a second exposure. Z-stacks with 2-3 planes and 7um separation were taken to ensure in-focus images were taken across the plate and 9 central fields of view were acquired per well which covered almost the entire well area.

#### Image Analysis: 2-color Cell-Cell fusion quantification

In the Perkin Elmer Harmony software, Maximum Projection, Brightfield correction, and Basic Flat Field Correction are used for analysis of each field of view. Nuclei are segmented using “Find Nuclei” function and Method C. Nuclei are selected for areas greater than 50μm^2^ (to eliminate debris). The GFP+ regions in each image are segmented into individual objects using an intensity threshold of 210-Infinity (manually identified to segment foreground from background). The area of GFP regions is calculated and regions with areas larger than 100μm^2^ are selected (this eliminates small regions/pixels that cross the intensity threshold; for comparison true syncytialized areas are generally tens of thousands of μm^2^). Similarly, the mCherry+ regions in each image are segmented into individual objects using an intensity threshold of 270-Infinity (manually identified to segment foreground from background). The area of mCherry regions is calculated and regions with areas larger than 100μm^2^ are selected. To avoid artificial regions of mCherry and GFP at nearby cell boundaries, all selected mCherry and GFP regions are resized by eroding the regions by 5 pixels. The pixel-based % overlap of selected Nuclei with the resized GFP+ regions is calculated using the Cross Population function. Similarly, the pixel-based % overlap of selected Nuclei with the resized mCherry+ regions is calculated using the Cross Population function. Nuclei are then filtered by using a Boolean Operator: nuclei having a GFP Nuclear overlap >90% AND a mCherry Nuclear overlap >90% are labeled as ++ Nuclei. Secondly, nuclei having a GFP Nuclear overlap >70% OR a mCherry Nuclear overlap >70% are labeled as Singly + Nuclei. The ratio of fused nuclei is then calculated as ++ Nuclei / Singly + Nuclei. Each datapoint is reported as the mean ratio of ++ Nuclei / Singly + Nuclei in a given well (averaging across all 9 fields of view). For each experiment, 6 technical replicates were imaged in separate wells and 3 biological replicates of each experiment were performed on different days.

#### SplitGFP Cell-cell fusion quantification

In the Perkin Elmer Harmony software, Maximum Projection, Brightfield correction, and Basic Flat Field Correction are used for analysis of each field of view. First, the GFP channel image is smoothed with a Gaussian filter, width = 2 pixels. GFP+ regions are segmented using an intensity threshold of 400-Infinity. The area of GFP regions is calculated and regions with areas larger than 100μm^2^ are selected to eliminate small debris. To account for differences in plating densities and the number of nuclei, the total DAPI+ region is segmented using an intensity threshold of 350-Infinity. The total area of each field of view is calculated using the Find Image Region, method= Whole Image Region. The area of the thresholded GFP region as well as the area of the Whole Image Region are calculated. The normalized GFP Area/DAPI Area is calculated as (GFP Area / Total Area) / ((DAPI Area / Total Area)/(Biological Replicate Mean(DAPI Area / Total Area)).

### Lysosome Imaging

#### Labeling

BeWo cells were plated at 10,000 cells per well into a Perkin Elmer CellCarrier Ultra 96-well plate. Cells were allowed to adhere 6 hours to overnight then treated with 0.1% DMSO or 20uM Forskolin for 48 hours. Media was changed every 24 hours. The media was then replaced with 100µLper well phenol-free supplemented DMEM with 1μg/mL Hoechst 33342 (Invitrogen) and 1:1000 Lysoview-540 (Biotium). Cells were incubated in this medium for 30min then imaged without washing.

#### Confocal Imaging

For Lysoview-540 imaging, a 40x water NA=1.1 objective was used and the following laser and filter sets were used to acquire three channels: Lysoview-540 was imaged with 561nm excitation, 570-630nm Emission, 500ms exposure at 100% power; Brightfield Transmission was imaged with Emission 650-760nm, 300ms exposure at 100% power; Hoechst was imaged with 375nm Excitation, 435-480nm Emission, 200ms exposure at 70% power. Hoechst/Brightfield were collected in the same exposure and separated from GFP which was collected in a second exposure. Five Z-stacks at 0.5um separation were acquired and 20 central fields of view were acquired per well which represents ∼ 1/5 of the well.

#### Image Analysis: Lysosomal Spots and Area

In the Perkin Elmer Harmony software, Individual Planes stack processing, Brightfield correction, and Advanced Flat Field Correction are used for analysis of each field of view. Since lysosomes could be seen moving during live acquisition of the relatively slow Z-stacks, for processing only one middle plane (Plane 2 of 5) was used for analysis rather than maximum projections. To identify the number of nuclei per image, the Hoechst channel was Gaussian smoothed with a filter of width = 8 pixels and Nuclei identified with the Find Nuclei function, Method C. Lysosomal spots were identified with the Find Spots function in the Lysoview-540nm (Cy3) channel using Method D. The Thresholded Lyso Region was identified using an absolute intensity threshold of 250-Infinity which was manually selected to separate foreground from background. The Lysoview-540 intensity and area were then calculated for each of these regions. The reported value (Thresholded Lyso Region Area Sum / # Nuclei) was calculated by summing the Thresholded Lyso Region Area per field of view and dividing by the total number of nuclei in that field. The reported value (# Lyso Spots / # Nuclei per well) was similarly calculated by dividing the total number of spots detected with Find Spots by the total number of nuclei detected per field of view. Each datapoint is reported as the mean value across all fields of view in a given well (averaging across all 20 fields of view). For each experiment, 6 technical replicates were imaged in separate wells and 3 biological replicates of each experiment were performed on different days.

#### Image Analysis: Nuclear Area Quantification

Using the three channel imaging data acquired for the lysosome imaging, the dataset was reprocessed to measure nuclear area in max projection Z-stacks. In the Perkin Elmer Harmony software, Maximum Projection, Brightfield correction, and Advanced Flat Field Correction are used for analysis of each field of view. Nuclei are segmented using the “Find Nuclei” function with Method C. Nuclei are selected for areas greater than 50μm^2^ (to eliminate debris) and border objects on the edge of each image are excluded. The area of each nucleus is then calculated. Each datapoint is reported as the mean Nuclear Area across all selected nuclei in a given well (averaging across all 20 fields of view). For each experiment, 6 technical replicates were imaged in separate wells and 3 biological replicates of each experiment were performed on different days.

### Nuclear/Cytoplasmic Halo-TFEB Imaging

#### Stable cell line

Halo-TFEB BeWo cells were created by plating 3E5 wild type BeWo cells per well into a 6-well plate and transfecting 0.8μg PiggyBac L30 promoter 3xF-Halo-TFEB IRES Puro along with 0.4μg Super Piggybac Transposase expressed on a second plasmid with Lipofectamine 3000 per the manufacturer’s instructions (Table 4). Cells were selected in 1μg/mL Puromycin for 2 weeks and maintained in Puromycin selection conditions.

#### Labeling

L30 Halo-TFEB BeWo cells were plated at 10,000 cells per well into a Perkin Elmer CellCarrier Ultra 96-well plate. Cells were allowed to adhere 6 hours to overnight then treated with 0.1% DMSO, 250nM Torin1 for 1 hour, or 20uM Forskolin for 24 or 48 hours. Media was changed every 24 hours. The media was then replaced with 100µLper well supplemented F12-K medium with 500nM JF646 Halo-tag ligand and 1:1000 MembraneSteady-488nm (Biotium).

Cells were incubated in this medium for 30min, then the medium was changed to phenol-free supplemented DMEM with 2μg/mL Hoechst 33342 (Invitrogen) + 1:1000 Enhancer (Biotium) and incubated for ∼10-15min before starting imaging. All medium (labeling, washes) contained the indicated drug conditions throughout and during imaging.

#### Confocal Imaging

For Halo-JF646 imaging, a 10x water NA=0.3 objective was used and the following laser and filter sets were used to acquire three channels: JF646 was imaged with 640nm excitation, 650-760nm emission for collection with 1500ms exposures at 100% power; Hoechst was imaged with 375nm Excitation, 435-480nm Emission, 400ms exposure at 100% power; and GFP was imaged with 488nm Excitation, 500-550 Emission, 600ms exposure at 100% power. GFP/JF646 were collected in the same exposure and separated from Hoechst which was collected in a second exposure. Three Z-stacks at 7.0um separation were acquired and 9 central fields of view were acquired per well which covered almost the entire well area.

#### Image Analysis: Nuclear/Cytoplasmic Quantification

In the Perkin Elmer Harmony software, Maximum Projection, Brightfield correction, and Basic Flat Field Correction are used for analysis of each field of view. Nuclei are segmented using the “Find Cells” function with Method C. Nuclei are selected for areas greater than 50μm^2^ (to eliminate debris) and border objects on the edge of each image are excluded. The cytoplasm is then segmented with the “Find Surrounding Region function with Method A. The mean JF646-channel intensity is then calculated within the Nuclei, Cytoplasm regions. Background is thresholded from all foreground JF646+ regions using an intensity cutoff of 200. The mean JF646-channel intensity is then calculated for the Background region. The background-corrected Nuclear/Cytoplasmic ratio is then calculated as (Mean Nuclear JF646 Intensity – Mean Background J6F646 Intensity) / (Mean Cytoplasmic JF646 Intensity – Mean Background J6F646 Intensity). Each datapoint is reported as the mean background-corrected Nuclear/Cytoplasmic ratio across all selected nuclei in a given well (averaging across all 9 fields of view). For each experiment, 5 technical replicates were imaged in separate wells and 3 biological replicates of each experiment were performed on different days.

### Single molecule tracking and analysis

#### Halo-H2B and Halo-NLS BeWo stable cell lines

Wild-type BeWo cells were transduced with concentrated L30 Halo-H2B IRES Neo or L30 Halo-NLS IRES Puro Lentivirus at an MOI of ∼0.5-1.0 (Table 4). Two days post transduction, cells were selected in 1μg/mL Puromycin (Halo-NLS) or 1600μg/mL Neomycin (G418) for 2 weeks and maintained in antibiotic selection conditions.

#### Sample preparation

BeWo cells stably L30 Halo-TFEB, Halo-H2B, or Halo-NLS were plated with 200,000 cells on Matek dishes containing No. 1.5 coverslip bottoms and let grow overnight. The next day, the media was exchanged for 20uM Forskolin or 0.1% DMSO-containing supplemented F12-K media and cells grown for 48 hours. All media was changed every 24hrs. For the 24hr Forskolin-treated samples, the sample was treated with 0.1% DMSO for the first 24hrs then changed into 20uM Forskolin at the 24hr media change timepoint. Before imaging, cells were stained simultaneously by changing the media to supplemented F12-K media containing 50nM JF549 and 25nM (for TFEB) or 5nM (for H2B and NLS) JFX646 in media containing Forskolin or DMSO, respectively. The slightly higher concentration JF549 was used to identify nuclei and for nuclear/cytoplasmic segmentation and JFX646 was used for tracking. Cells were stained with the indicated JF dyes in the incubator for 30 minutes, then rinsed with PBS, then washed in supplemented F-12K media containing 20uM Forskolin or 0.1% DMSO respectively for at least 30 min. Cells were left in this wash media until just before imaging. Just prior to imaging, cells were again rinsed with PBS then put into phenol-free high-glucose DMEM media containing 10% FBS and supplemented with Sodium Pyruvate and Glutamax with 0.1% DMSO or 20uM Forskolin respectively.

#### Acquisition

Single molecule tracking was performed as previously described by Hansen et al. *Elife* 2017 with some minor modifications^65^. Single-molecule imaging was performed on a custom-built Nikon TI microscope with a 100x/NA 1.49 oil-immersion TIRF objective (Nikon apochromat CFI Apo TIRF 100x Oil), EM-CCD camera (Andor, Concord, MA, iXon Ultra 897; frame-transfer mode; vertical shift speed: 0.9 μs; −70°C), a perfect focusing system to correct for axial drift and motorized laser illumination (Ti-TIRF, Nikon), to achieve highly inclined and laminated optical sheet illumination. The incubation chamber maintained a humidified 37°C atmosphere with 5% CO_2_ and the objective was also heated to 37°C. Excitation was achieved using the following laser lines: 561 nm (1 W, Genesis Coherent, Santa Clara, CA) for JF549; 633 nm (1 W, Genesis Coherent, Pala Alto, CA) for JFX646; 405 nm (140 mW, OBIS, Coherent) for all experiments. The excitation lasers were modulated by an acousto-optic Tunable Filter (AA Opto-Electronic, France, AOTFnC-VIS-TN) and triggered with the camera TTL exposure output signal. The laser light is coupled into the microscope by an optical fiber and then reflected using a multi-band dichroic (405 nm/488 nm/561 nm/633 nm quad-band, Semrock, Rochester, NY) and then focused in the back focal plane of the objective. Fluorescence emission light was filtered using a single band-pass filter placed in front of the camera using the following filters: JF549: Semrock 593/40 nm bandpass filter; JFX646: Semrock 676/37 nm bandpass filter. The microscope, cameras, and hardware were controlled through NIS-Elements software (Nikon). The pixel size in this configuration is 160nm.

For TFEB masking, prior to each movie, a single exposure of ∼20-100ms 561 nm excitation (10-20% AOTF) was used to record the JF549 fluorescence signal as cytoplasmic or nuclear within the specified ROI used to subsequently record the SMT movies. Given the variable expression levels between cells, the exposure time and illumination power for this mask exposure were sometimes adjusted manually to ensure the cell intensities were not outside the linear range of the camera. Movies were acquired with stroboscopic illumination and dark state reactivation of fluorophores during the camera integration time as follows: 1 ms 633 nm excitation (100% AOTF) of JFX646 was delivered at the beginning of the frame and 405 nm photo-activation pulses (10% AOTF) were delivered during the camera integration time (∼447 μs) to achieve a desired mean reactivation of ∼1-10 molecules per frame. 8,000 frames were recorded per cell per experiment. 150-pixel by 150-pixel ROIs were user-selected to be centered on a nucleus (identified via JF549 signal) and were acquired per cell with a camera exposure time of 7 ms. Around 20 cells were recorded for each condition and each condition was performed in triplicate on separate days for three biological replicates.

#### Localization and Tracking Analysis with quot

Localization and tracking of molecules were performed using the ‘quot’ package (available at https://github.com/alecheckert/quot). While tracking was performed over almost the entire movie (starting at frame 100), very conservative tracking parameters were used to ensure that high density at the beginning of the movie did not result in many misconnected trajectories. Specifically, method = ‘conservative’, and max_spots_per_frame = 7 were used for this purpose which will exclude more dense localizations and any ambiguous connections from contributing to valid trajectories. The following configuration settings were specified in the quot config.toml file: [filter] start = 100, method = ‘identity’, chunk_size = 100; [detect] method = ‘llr’, k = 1.2, w = 15, t = 18.0; [localize] method = ‘ls_int_gaussian’, window_size = 9, sigma = 1.0, ridge = 0.001, max_iter = 20, damp = 0.3; [track] method = ‘conservative’, max_spots_per_frame = 7, pixel_size_um = 0.16, frame_interval = 0.00748, search_radius = 1.0, max_blinks = 0, min_I0 = 0.0, scale = 7.0.

#### Bayesian Inference of Brownian Diffusion using saSPT

Modeling of diffusion from trajectories was performed using the Bayesian state array approach encoded in the ‘saspt’ package (available at https://github.com/alecheckert/saspt). To avoid dense localizations or potential misconnected trajectories at the beginning of the movie biasing diffusion states, analysis was excluded for the first 1000 frames of each movie. The StateArray class (from saspt.sa) was used for diffusion analysis and the following configuration settings were specified: start_frame = 1000, pixel_size_um = 0.16, frame_interval = 0.00748, focal_depth = 0.7, sample_size = 1000000, likelihood = ‘rbme’, splitsize = 10. For subsampling the trajectories to the lowest number of any dataset, the sample_size was set to 20,000 instead. The “bound fraction” was defined as the cumulative posterior occupation for diffusion coefficients less than 0.1 μ^2^/s. Movies were collected over three independent days of imaging with 15-20 cells collected per condition per day.

#### Nuclear Masking

Nuclear masks were created in FIJI using the single JF549 exposures collected prior to each movie. For each image, the following processing was performed using an automated batch script in FIJI: run(“Gaussian Blur…”, “sigma=2”); setAutoThreshold(“Minimum dark”); run(“Convert to Mask”); run(“Fill Holes”); run(“Options…”, “iterations=2 count=1 black pad do=Erode”). Masks were then adjusted by hand in FIJI to ensure that a single nucleus was correctly masked in each image. For example, if another overlapping nearby nucleus was captured in the JF549 exposure, this region of the mask was erased to only include one nucleus per mask. For dim images where the automated threshold setting did not perform well, nuclear regions were drawn manually, binarized to create the nuclear masks, then holes filled and eroded as above. For cells where the TFEB signal was predominantly cytoplasmic, images were inverted before running the process above or nuclear regions were identified manually.

### 3xFlag-Halo-TFEB Chromatin Immunoprecipitation (ChIP) and Analysis

#### Cell sample preparation and Immunoprecipitation

Stably expressing L30 3xFlag-Halo-TFEB IRES Puro BeWo cells (passage ∼10) were scaled up to two 15cm plates per condition. Cells were grown to ∼50% confluence, then media was changed to 0.1% DMSO or 20uM Forskolin and treated for 48 hours, changing media at 24 hours. For Torin1 treated samples, cells were treated with 0.1% DMSO for 46 hours, then changed to media containing 250nM Torin1 for 2 hours before being fixed. After treatment, media on each 15cm plate was changed to 20mL of serum-free F12-K media containing 1% formaldehyde and incubated shaking for 5 minutes. Crosslinking was then quenched by adding 10mL of 0.125M glycine in PBS and shaking for 5 additional minutes. Media was then poured off and plates rinsed twice with ice cold PBS. Finally, 5mL of PBS plus protease inhibitors (0.25uM PMSF, 10ug/mL aprotinin) was added to each plate and cells were scraped on ice, collected in a 15mL conical, then centrifuged at 1200rpm for 10 minutes at 4°C. The supernatant was aspirated and cell pellets were flash frozen in liquid nitrogen and stored at -80°C until further processing. Plating, treatment, and fixation of cells was repeated twice for two biological replicates of each condition.

Cell pellets were thawed on ice, resuspended in 2 ml of cell lysis buffer (5 mM PIPES, pH 8.0, 85 mM KCl, and 0.5% NP-40; 1ml/15 cm plate) w/ protease inhibitors and incubated for 10 minutes on ice. During incubation, the lysates were pipetted up and down every 5 minutes. Lysates were then centrifuged for 10 minutes at 4000 rpm. Nuclear pellets were resuspended in 6 volumes of sonication buffer (50 mM Tris-HCl, pH 8.1, 10 mM EDTA, 0.1% SDS), w/ protease inhibitors, incubated on ice and, sonicated to obtain DNA fragments below 1000 bp in length (Covaris S220 sonicator, 20% Duty factor, 200 cycles/burst, 150 peak incident power, 6-8 cycles 30 seconds on and 30 seconds off). Sonicated lysates were cleared by centrifugation and chromatin (400 ug per antibody) was diluted in RIPA buffer (10 mM Tris-HCl, pH 8.0, 1 mM EDTA, 0.5 mM EGTA, 1% Triton X-1000, 0.1% SDS, 0.1% Na-deoxycholate, 140 mM NaCl) to a final concentration of 0.8 mg/ml, precleared with magnetic Protein G Dynabeads (Thermo Fisher, #10009D) for 2 hours at 4°C. Precleared lysates were then immunoprecipitated overnight with 4 mg of normal anti-mouse IgG (Jackson ImmunoResearch, #211-032-171) and anti-FLAG [M2] (Sigma Aldrich, # F1804). About 4% of the precleared chromatin was saved as input. Immunoprecipitated DNA was purified with Qiagen QIAquick PCR Purification Kit (Qiagen, #28106), eluted in 40uL of 0.1x TE (1 mM Tris-HCl pH 8.0, 0.01 mM EDTA). Samples were checked by qPCR together with 2% of the input chromatin then used in ChIP-seq library preparation.

#### Library preparation

ChIP-seq libraries were prepared independently from two ChIP biological replicates using the NEBNext Ultra II DNA Library Prep Kit (New England Biolabs, #E6177L) according to manufacturer’s instructions with a few modifications. As starting material, 20ng of ChIP input DNA (as measured by Nanodrop) and 20uL of the immunoprecipitated DNA, spiked with 5uL of sheared *Drosophila melanogaster* DNA (10 ng/mL) were used. The recommended reagents’ volumes were cut in half. The NEBNext Adaptor for Illumina was diluted 1:10 in Tris/NaCl, pH 8.0 (10 mM Tris-HCl pH 8.0, 10 mM NaCl) and the ligation step extended to 30 minutes. A single purification step with 0.9x volumes of Agencourt AMPure XP PCR burification beads (Beckman Coulter, # A63880) was performed, after ligation. DNA was eluted in 22uL of 10 mM

Tris-HCl (pH 8.0), and 20uL of the eluted DNA was used for library enrichment step. Library samples were enriched with 11 cycles of PCR amplification in 50uL of total reaction volume (10uL 5x KAPA buffer, 1.5uL 10 mM dNTPs, 0.5uL 10 mM NEB Universal PCR primer, 0.5uL of 10 mM index primers, 1 mL KAPA polymerase, 16.5uL nuclease-free water and 20uL sample). After amplification, PCR samples were again purified with 0.9x volumes of AMPure XP PCR amplification beads and eluted in 33uL of 10 mM Tris-HCl (pH 8.0). Library concentration was assessed using Qubit dsDNA HS Assay Kit (Invitrogen, #Q32851). Libraries were sent to Medgenome Inc (Foster City, CA) for fragment analysis, multiplexing and sequencing on Illumina Novaseq 6000 platform (150 bp, paired end reads). Nine multiplexed libraries (Input, mFlag, IgG in DMSO, Torin and Forskolin treated samples) were pooled and sequenced per lane.

#### ChIP-seq analysis

ChIP-seq raw reads from FLAG-Halo-TFEB BeWo cells treated with DMSO, Torin and Forskolin were quality checked with FastQC (Version 0.10.1) and aligned to the human genome (hg38 assembly) using Bowtie2 (version 2.3.4.1) with options: --local --very-sensitive-local --no-unal --no-mixed --no-discordant -p10 -I 50 -X 1000. Samtools^66^ (version 1.8) were used to create, sort and index bam files. Peaks were called with MACS2, version 2.1.0.20140616 (-f BAMPE --nomodel) using either the input or IgG DNA as controls. Similar numbers of peaks were called against each control, so the peaks called against the IgG sample were used in further analysis.

Heatmaps were created using deepTools (Version 2.4.1).^67^ First, bam files were converted to bigWig files with normalized read numbers to 1x sequencing depth, obtaining read coverage per 50 bp bins across the whole genome, using bamCoverage (-of bigwig --binSize 50 – normalizeTo1x 2913022398 --extendReads --ignoreDuplicates). To perform IDR analysis, MACS2 was used to call peaks with a relaxed cutoff (macs2 callpeak -B -p 1e-3 --nomodel -B -p 1e-3 -t), peaks were then sorted by their p-value and the sorted narrowPeak files were used as inputs for the IDR program. Significant peaks were identified as those with a transformed IDR value >=540, which is equivalent to a p-value of <=0.05. For differential enrichment analysis, MACS2 peak calling was repeated with higher stringency but a fixed fragment size (macs2 callpeak -B -t -f BAM -g hs --nomodel --extsize 194). The fixed fragment size was calculated by the average of the predicted fragment length of the FSK FLAG and Torin FLAG sorted bam files (using the command macs2 predictd -i). Differentially enriched peaks were identified by running -g macs2 bdgdiff (options -g 60 -l 120) and the sequencing depths for each condition were identified from the “tags after filtering in control” from the MACS2 callpeak output. Deeptools was used to create heatmaps of these differentially enriched peaks using the normalized bamCoverage bigWig files to compute reads across 6kB centered on TFEB peak and sorted by decreasing TFEB (FLAG) enrichment. Finally, differentially-enriched peaks identified by bdgdiff analysis and statistically robust peaks identified by IDR analysis were overlapped using bedtools intersect and the resulting bed files were mapped to nearby annotated genomic regions using ChIPSeeker with a binding region of -2000bp to +500bp using annotated data from TxDb.Hsapiens.UCSC.hg38.knownGene and enriched GO categories and dotplots were made with ClusterProfiler.^68,69^ Summits from these differentially-enriched (MACS2 bdgdiff) and statistically robust (IDR p-value <0.05) were derived from MACS2, expanded 250bp on either side (using bedtools slop function in Galaxy), filtered for unique entries (using FASTA Merge Files and Filter Unique Sequences function in Galaxy) then input into MEME-ChIP (online tool, Version 5.5.5) for motif discovery in the HOCOMOCO Human (v11 FULL) database.^49,50^

## Statistics and plotting

Western blot images were prepared using FIJI and Adobe Illustrator. RNASeq plots were made using ggplot2 in R with Jupyter notebook. Statistical tests and bar charts were made in GraphPad Prism version 10. Genomics track displays were created using IGV viewer (version 2.16.2) and Adobe Illustrator. GO term analysis was performed at https://geneontology.org/.

## Acknowledgements

MNE was financially supported by the HS Chau Foundation and the NIH Stem Cell Fellowship T32GM098218. JM was supported by the NIH training grant T32GM139780. We thank Gina Dailey who provided some of the plasmids used in this study and Claudia Cattoglio who provided Lentivirus reagents and advice on the ChIP sequencing analysis workflow. We acknowledge Gokul Upadhyayula who provided useful advice on cell segmentation approaches for quantification. We thank Fyodor Urnov for helpful advice on CRISPR editing with RNPs. We thank Roberto Zoncu who provided the TFEB cDNA plasmid and useful advice for this project. We thank Mary West and the QB3 High-Throughput Screening Facility (HTSF) at UC Berkeley. This work was performed in part in the QB3 HTSF that provided the Opera Phenix microscope, supported by the Office of the Director, National Institutes of Health, under Award Number S10OD021828. The content is solely the responsibility of the authors and does not necessarily represent the official views of the National Institutes of Health.

## Author Contributions

MNE conceptualized the project, developed methodology, performed most experiments, performed analysis, visualized results, wrote the original draft and edited the manuscript. LD performed ChIP experiments and edited the manuscript. VF wrote software used in the analysis of SMT data. JM performed preliminary optimization for ChIP. EY supported characterization of cell lines and cloning. RT and XD supervised the project, supported funding of the project, and edited the manuscript.

## References

1. Hubrecht, A. A. W. Memoirs: Studies in Mammalian Embryology: I.--The Placentation of Erinaceus Europæus, with Remarks on the Phylogency of the Placenta. J Cell Sci s2–30, 283–404 (1889).

2. Burton, G. J. & Fowden, A. L. The placenta: a multifaceted, transient organ. Philosophical Transactions of the Royal Society B: Biological Sciences 370, 20140066 (2015).

3. Turbeville, H. R. & Sasser, J. M. Preeclampsia beyond pregnancy: long-term consequences for mother and child. 10.1152/ajprenal.00071.2020 318, F1315–F1326 (2020).

4. Brosens, I., Pijnenborg, R., Vercruysse, L. & Romero, R. The “Great Obstetrical Syndromes” are associated with disorders of deep placentation. Am J Obstet Gynecol 204, 193–201 (2011).

5. 5. Davis, N., Smoots, A. & Goodman, D. Pregnancy-Related Deaths: Data from 14 U.S. Maternal Mortality Review Committees, 2008-2017. Centers for Disease Control and Prevention, U.S. Department of Health and Human Services https://www.cdc.gov/reproductivehealth/maternal-mortality/erase-mm/mmr-data-brief.html (2019).

6. Ghulmiyyah, L. & Sibai, B. Maternal Mortality From Preeclampsia/Eclampsia. Semin Perinatol 36, 56–59 (2012).

7. Turco, M. Y. & Moffett, A. Development of the human placenta. Development (Cambridge) 146, (2019).

8. Vargas, A. et al. Reduced expression of both syncytin 1 and syncytin 2 correlates with severity of preeclampsia. Reproductive Sciences 18, 1085–1091 (2011).

9. Mukherjee, I. et al. Oxidative stress-induced impairment of trophoblast function causes preeclampsia through the unfolded protein response pathway. Scientific Reports 2021 11:1 11, 1–20 (2021).

10. Shao, X. et al. Placental trophoblast syncytialization potentiates macropinocytosis via mTOR signaling to adapt to reduced amino acid supply. Proc Natl Acad Sci U S A 118, (2021).

11. Galton, M. DNA content of placental nuclei. J Cell Biol 13, (1962).

12. Pierce, G. B. & Midgley, A. R. THE ORIGIN AND FUNCTION OF HUMAN SYNCYTIOTROPHOBLASTIC GIANT CELLS. Am J Pathol 43, 153–73 (1963).

13. Ellery, P. M., Cindrova-Davies, T., Jauniaux, E., Ferguson-Smith, A. C. & Burton, G. J. Evidence for Transcriptional Activity in the Syncytiotrophoblast of the Human Placenta. Placenta 30, 329 (2009).

14. Sha, M. et al. Syncytin is a captive retroviral envelope protein involved in human placental morphogenesis. Nature 2000 403:6771 403, 785–789 (2000).

15. Blond, J.-L. et al. An Envelope Glycoprotein of the Human Endogenous Retrovirus HERV-W Is Expressed in the Human Placenta and Fuses Cells Expressing the Type D Mammalian Retrovirus Receptor. J Virol 74, 3321–3329 (2000).

16. Blaise, S., De Parseval, N., Bénit, L. & Heidmann, T. Genomewide screening for fusogenic human endogenous retrovirus envelopes identifies syncytin 2, a gene conserved on primate evolution. Proc Natl Acad Sci U S A 100, 13013–13018 (2003).

17. Roberts, R. M. et al. Syncytins expressed in human placental trophoblast. Placenta 113, 8–14 (2021).

18. Esnault, C. et al. A placenta-specific receptor for the fusogenic, endogenous retrovirus-derived, human syncytin-2. Proc Natl Acad Sci U S A 105, (2008).

19. Toufaily, C. et al. MFSD2a, the Syncytin-2 receptor, is important for trophoblast fusion. Placenta 34, 85–88 (2013).

20. Papuchova, H. & Latos, P. A. Transcription factor networks in trophoblast development. Cellular and Molecular Life Sciences 2022 79:6 79, 1–17 (2022).

21. Lin, C., Lin, M. & Chen, H. Biochemical characterization of the human placental transcription factor GCMa/1. Biochemistry and Cell Biology 83, (2005).

22. Yu, C. et al. GCMa Regulates the Syncytin-mediated Trophoblastic Fusion. Journal of Biological Chemistry 277, 50062–50068 (2002).

23. Liang, C. Y. et al. GCM1 Regulation of the Expression of Syncytin 2 and Its Cognate Receptor MFSD2A in Human Placenta. Biol Reprod 83, 387–395 (2010).

24. Schreiber, J. et al. Placental Failure in Mice Lacking the Mammalian Homolog of Glial Cells Missing, GCMa. Mol Cell Biol 20, (2000).

25. Simmons, D. G. et al. Early patterning of the chorion leads to the trilaminar trophoblast cell structure in the placental labyrinth. Development 135, 2083–2091 (2008).

26. Bainbridge, S. A. et al. Effects of reduced Gcm1 expression on trophoblast morphology, fetoplacental vascularity, and pregnancy outcomes in mice. Hypertension 59, (2012).

27. Anson-Cartwright, L. et al. The glial cells missing-1 protein is essential for branching morphogenesis in the chorioallantoic placenta. Nature Genetics 2000 25:3 25, 311–314 (2000).

28. Baczyk, D. et al. Glial cell missing-1 transcription factor is required for the differentiation of the human trophoblast. Cell Death & Differentiation 2009 16:5 16, 719–727 (2009).

29. Jeyarajah, M. J. et al. The multifaceted role of GCM1 during trophoblast differentiation in the human placenta. Proc Natl Acad Sci U S A 119, e2203071119 (2022).

30. Steingrímsson, E., Tessarollo, L., Reid, S. W., Jenkins, N. A. & Copeland, N. G. The bHLH-Zip transcription factor Tfeb is essential for placental vascularization. Development 125, 4607–4616 (1998).

31. Fisher, D. E., Carr, C. S., Parent, L. A. & Sharp, P. A. TFEB has DNA-binding and oligomerization properties of a unique helix-loop-helix/leucine-zipper family. Genes Dev 5, 2342–2352 (1991).

32. Tan, A., Prasad, R., Lee, C. & Jho, E. hoon. Past, present, and future perspectives of transcription factor EB (TFEB): mechanisms of regulation and association with disease. *Cell* Death & Differentiation 2022 29:8 29, 1433–1449 (2022).

33. Palmieri, M. et al. Characterization of the CLEAR network reveals an integrated control of cellular clearance pathways. Hum Mol Genet 20, (2011).

34. Sardiello, M. et al. A gene network regulating lysosomal biogenesis and function. Science (1979) 325, 473–477 (2009).

35. Puertollano, R., Ferguson, S. M., Brugarolas, J. & Ballabio, A. The complex relationship between TFEB transcription factor phosphorylation and subcellular localization. EMBO J 37, e98804 (2018).

36. Roczniak-Ferguson, A. et al. The transcription factor TFEB links mTORC1 signaling to transcriptional control of lysosome homeostasis. Sci Signal 5, (2012).

37. Settembre, C. et al. A lysosome-to-nucleus signalling mechanism senses and regulates the lysosome via mTOR and TFEB. EMBO J 31, 1095–1108 (2012).

38. Nakashima, A. et al. Evidence for lysosomal biogenesis proteome defect and impaired autophagy in preeclampsia. Autophagy 16, 1771–1785 (2020).

39. Nakashima, A. et al. Placental autophagy failure: A risk factor for preeclampsia. Journal of Obstetrics and Gynaecology Research 46, 2497–2504 (2020).

40. Furuta, A. et al. The Autophagy-Lysosomal Machinery Enhances Cytotrophoblast-Syncytiotrophoblast Fusion Process. Reproductive Medicine 2022*, Vol.* 3, *Pages 112-126* **3**, 112–126 (2022).

41. Shao, X. et al. Placental trophoblast syncytialization potentiates macropinocytosis via mTOR signaling to adapt to reduced amino acid supply. Proc Natl Acad Sci U S A 118, (2021).

42. Pattillo, R. A. & Gey, G. O. The Establishment of a Cell Line of Human Hormone-synthesizing Trophoblastic Cells in Vitro. Cancer Res 28, 1231–1236 (1968).

43. Wice, B., Menton, D., Geuze, H. & Schwartz, A. L. Modulators of cyclic AMP metabolism induce syncytiotrophoblast formation in vitro. Exp Cell Res 186, 306–316 (1990).

44. Orendi, K., Gauster, M., Moser, G., Meiri, H. & Huppertz, B. The choriocarcinoma cell line BeWo: syncytial fusion and expression of syncytium-specific proteins. Reproduction 140, 759–766 (2010).

45. Hudon Thibeault, A. A., Vaillancourt, C. & Sanderson, J. T. Profile of CYP19A1 mRNA expression and aromatase activity during syncytialization of primary human villous trophoblast cells at term. Biochimie 148, (2018).

46. Martina, J. A., Chen, Y., Gucek, M. & Puertollano, R. MTORC1 functions as a transcriptional regulator of autophagy by preventing nuclear transport of TFEB. Autophagy 8, (2012).

47. Napolitano, G. et al. mTOR-dependent phosphorylation controls TFEB nuclear export. Nat Commun 9, 1–10 (2018).

48. Li, Q., Brown, J. B., Huang, H. & Bickel, P. J. Measuring reproducibility of high-throughput experiments. Annals of Applied Statistics 5, (2011).

49. Machanick, P. & Bailey, T. L. MEME-ChIP: Motif analysis of large DNA datasets. Bioinformatics 27, (2011).

50. Gaspar, J. M. Improved peak-calling with MACS2. bioRxiv (2018).

51. Steingrímsson, E. et al. Mitf and Tfe3, two members of the Mitf-Tfe family of bHLH-Zip transcription factors, have important but functionally redundant roles in osteoclast development. Proc Natl Acad Sci U S A 99, 4477 (2002).

52. Shimizu, T. et al. CRISPR screening in human trophoblast stem cells reveals both shared and distinct aspects of human and mouse placental development. Proceedings of the National Academy of Sciences 120, e2311372120 (2023).

53. Gerri, C. et al. Initiation of a conserved trophectoderm program in human, cow and mouse embryos. Nature 2020 587:7834 587, 443–447 (2020).

54. Nakashima, A. et al. Placental autophagy failure: A risk factor for preeclampsia. Journal of Obstetrics and Gynaecology Research 46, 2497–2504 (2020).

55. Oh, S.-Y. & Roh, C.-R. Autophagy in the placenta. Obstet Gynecol Sci 60, 241–259 (2017).

56. Tan, A., Prasad, R. & Jho, E. hoon. TFEB regulates pluripotency transcriptional network in mouse embryonic stem cells independent of autophagy–lysosomal biogenesis. Cell Death Dis 12, (2021).

57. Iyer, H., Shen, K., Meireles, A. M. & Talbot, W. S. Repression of lysosomal transcription factors Tfeb and Tfe3 is essential for the migration and function of microglia. bioRxiv 2020.10.27.357897 (2022) doi:10.1101/2020.10.27.357897.

58. Kim, S. et al. PARsylated transcription factor EB (TFEB) regulates the expression of a subset of Wnt target genes by forming a complex with β-catenin-TCF/LEF1. Cell Death & Differentiation 2021 28:9 28, 2555–2570 (2021).

59. Akiyama, Y., Hosoya, T., Poole, A. M. & Hotta, Y. The gcm-motif: A novel DNA-binding motif conserved in Drosophila and mammals (DNA-binding activit^y͞^ transcriptional regulator). Neurobiology 93, (1996).

60. Nait-Oumesmar, B., Copperman, A. B. & Lazzarini, R. A. Placental expression and chromosomal localization of the human Gcm 1 gene. Journal of Histochemistry and Cytochemistry 48, (2000).

61. Pan, Y. J. et al. Expression of urotensin II is associated with placental autophagy in patients with severe preeclampsia. Journal of Human Hypertension 2018 32:11 32, 759–769 (2018).

62. Oh, S. Y. et al. Autophagy-related proteins, LC3 and beclin-1, in placentas from pregnancies complicated by preeclampsia. Reproductive Sciences 15, 912–920 (2008).

63. Akaishi, R. et al. Autophagy in the placenta of women with hypertensive disorders in pregnancy. Placenta 35, 974–980 (2014).

64. Nakashima, A. et al. Impaired autophagy by soluble endoglin, under physiological hypoxia in early pregnant period, is involved in poor placentation in preeclampsia. Autophagy 9, 303–316 (2013).

65. Hansen, A. S., Pustova, I., Cattoglio, C., Tjian, R. & Darzacq, X. CTCF and cohesin regulate chromatin loop stability with distinct dynamics. Elife (2017) doi:10.7554/elife.25776.

66. Li, H. et al. The Sequence Alignment/Map format and SAMtools. Bioinformatics 25, (2009).

67. Ramírez, F. et al. deepTools2: a next generation web server for deep-sequencing data analysis. Nucleic Acids Res 44, (2016).

68. Yu, G., Wang, L. G. & He, Q. Y. ChIP seeker: An R/Bioconductor package for ChIP peak annotation, comparison and visualization. Bioinformatics 31, (2015).

69. Yu, G., Wang, L. G., Han, Y. & He, Q. Y. ClusterProfiler: An R package for comparing biological themes among gene clusters. OMICS 16, (2012).

