## Supplemental Information for "TFEB controls syncytiotrophoblast differentiation"

**A**

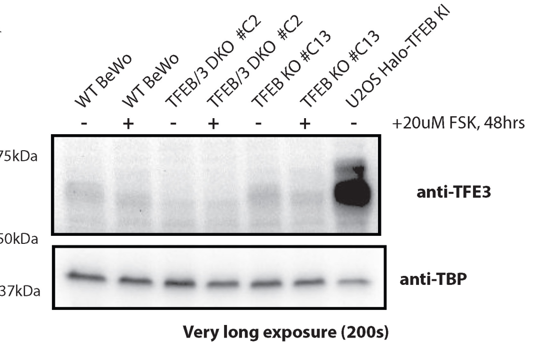


**B**

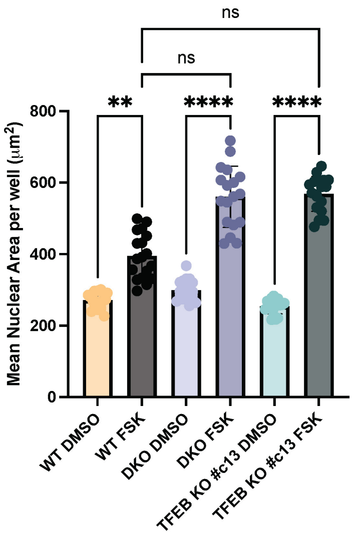


**Figure S1. Further assessment of TFEB/TFE3 perturbation in BeWo cells. A) Long exposure of the same TFE3** western blotting in Figure 1C in wild-type and CRISPR KO BeWo cells, and in U2-OS cells. **B)** Quantification of high-throughput confocal imaging measuring the mean per-well nuclear area for wild-type, DKO, and TFEB KO BeWo cells treated with DMSO or Forskolin for 48hrs. Statistical significance from Kruskal-Wallis ANOVA test with Dunn's multiple comparison test is shown where ns = not significant, * = p<0.05, ** = p<0.01, *** = p<0.001, **** and p<0.0001.

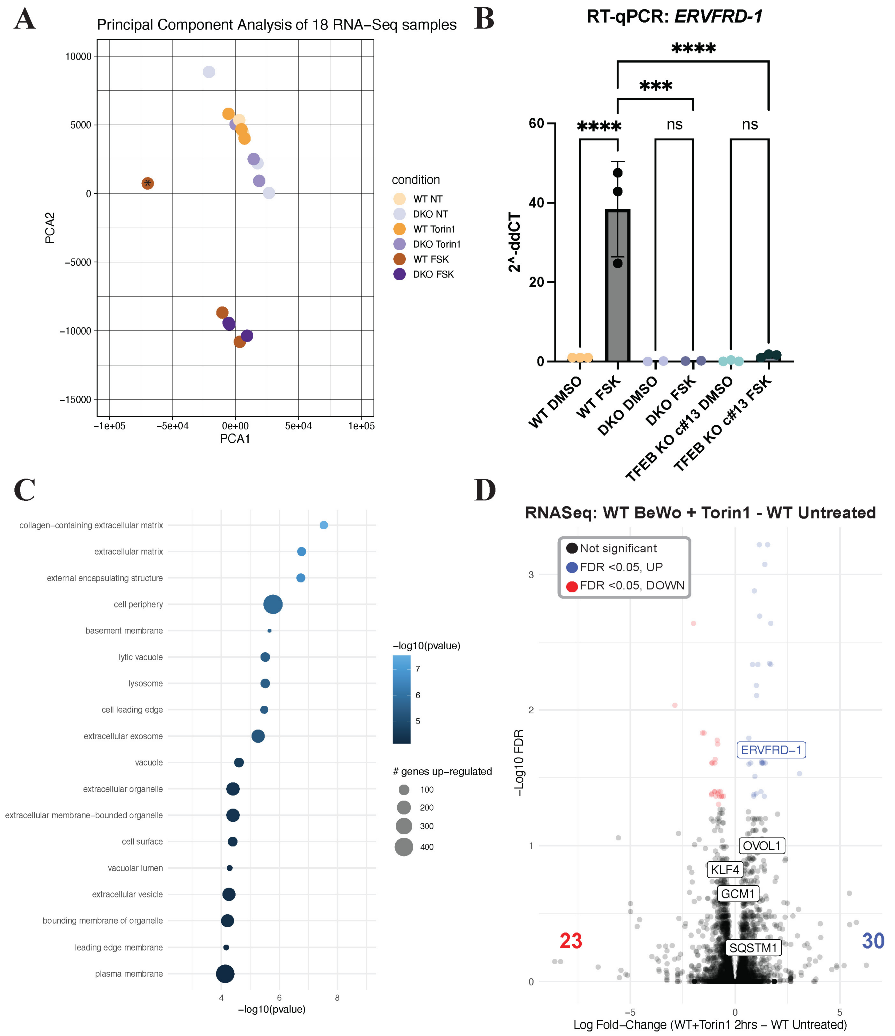


**Figure S2. Further assessment of RNASeq in TFEB/TFE3 DKO BeWo cells. A)** Principal Component Analysis of the six sample conditions with three replicates each. RNA-seq replicates show clustering of Forskolin-treated samples and separation from DMSO-treated and Torin-1 treated samples. One outlier sample, marked with an asterisk (*), was excluded from further analysis. **B)** Expression of *ERVFRD-1* transcripts measured by RT-qPCR and quantified by the delta-delta Ct method. Statistical significance from an ordinary one-way ANOVA with Tukey's multiple comparisons test is shown where ns = not significant, * = p<0.05, ** = p<0.01, *** = p<0.001, and **** = p<0.0001. **C)** Top 10 enriched GO Terms for Cellular Compartment of genes which are upregulated in the wild-type Forskolin-treated BeWos compared to the Forskolin-treated TFEB/TFE3 DKO cells. **D)** Volcano plot of RNA-Seq data comparing 2-hour Torin1-treated wild-type BeWo cells with 2-hour Torin1-treated TFEB/TFE3 DKO BeWo cells. Genes significantly upregulated in the Torin1-treated DKO cells are shown in red, and genes significantly downregulated in the Torin1-treated DKO cells are shown in blue.

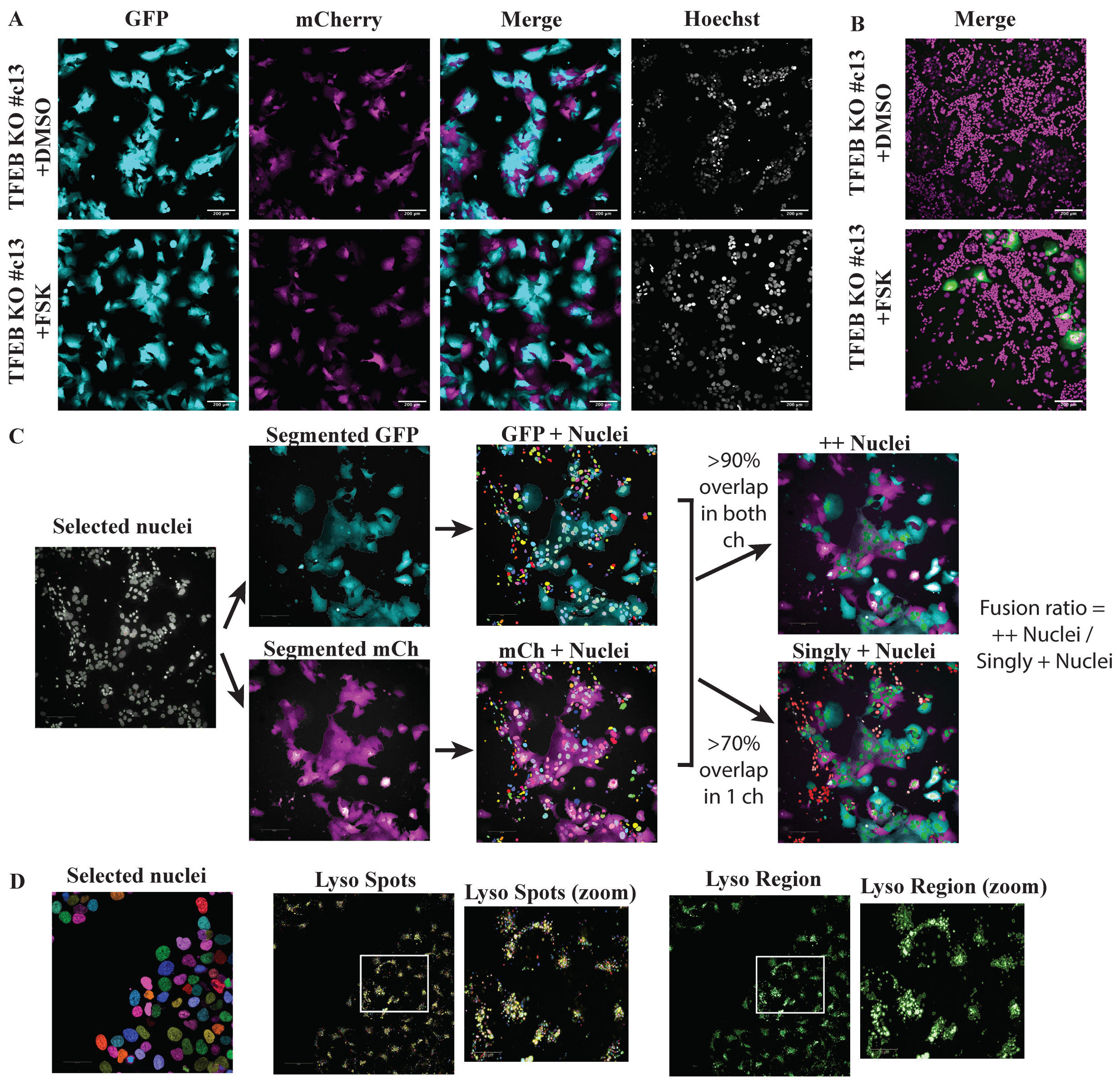


**Figure S3. TFEB KO cell-cell fusion defects and segmentation analysis pipelines. A)** Two color cell-cell fusion experiments co-culturing two populations of mCherry- and GFP-expressing TFEB KO #c13 BeWo cells were imaged on a spinning disk confocal microscope. Individual channels are shown in grayscale. In the merged composite image, the GFP channel is represented in cyan and the mCherry channel is represented in magenta. White arrows indicate fused syncytial areas in the Forskolin-treated wild-type cells. Scale bar = 200µm. **B)** Split-GFP cell-cell fusion experiments co-culturing GFP1-10 expressing TFEB KO #c13 BeWo cells with GFP11-expressing 293T cells were imaged on a spinning disk confocal microscope. In the merged composite image, the GFP channel is represented in green and the Hoechst channel is represented in magenta. Fused syncytial areas are shown by the reconstitution of GFP fluorescence shown in green. Scale bar = 200µm. Quantification of B and C are shown in Fig. 3C,D. **C)** Overview and example of the two color cell-cell fusion segmentation and quantification pipeline. Selected nuclei are shown in green outline. Subsequently, GFP (cyan) and mCherry (magenta) channels are separately thresholded and filtered to yield single channel-positive regions. Nuclei (rainbow) are overlaid on each channel and those with >90% overlap in both channel regions are deemed ++ (top right, green nuclei) while those with >70% overlap are defined as singly + (bottom right, green nuclei; excluded nuclei which do not meet this criteria are shown in red). In panel C only, images were adjusted for easy visualization of segmentation and gamma = 2.0 applied. **D)** Example of segmentation of live lysosomal imaging. Selected nuclei (left, rainbow) are used segmented and the counts used for normalization of cell density. Lysoview-540 imaged in the Cy3 channel (yellow) are segmented with a spot detector (rainbow). A close-up view of the segmentation quality is shown (yellow, right). To quantify the total area occupied by lysosomes, the Lysoview-540-positive region is thresholded and filtered (green) and the area summed. A close-up view of the segmentation quality is shown (green, far right).

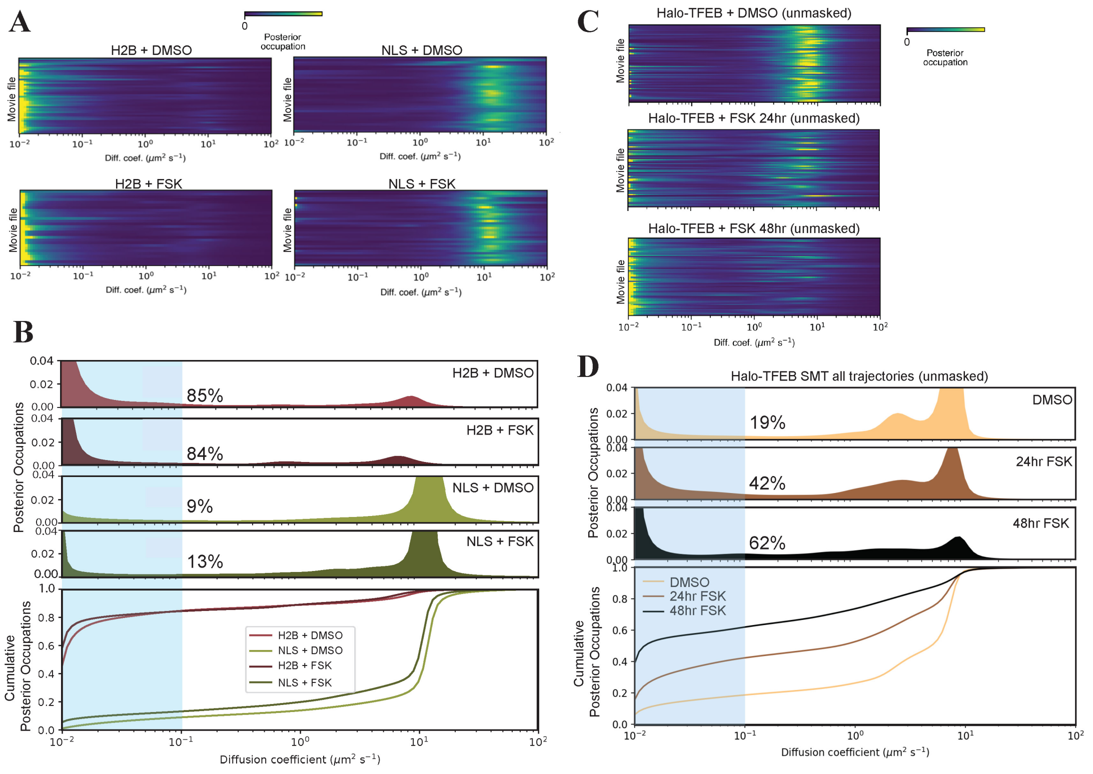


**Figure S4. Extended single molecule imaging data: controls and unmasked trajectories. A)** Heatmaps of diffusion coefficients following Bayesian analysis of single molecule trajectory data of control BeWo cells expressing Halo-H2B or Halo-NLS treated with DMSO or Forskolin for 48hrs. Each row corresponds to the distribution of posterior occupations for a single movie file. **B)** The distribution of diffusion coefficient occupancies for trajectories of Halo-NLS and Halo-H2B treated with DMSO or Forskolin for 48hrs. The fraction bound (calculated by the fraction of the distribution with a diffusion coefficient <0.1$\mu m^2/s$) is shown highlighted with the blue region and the quantified values displayed as black percentages. The cumulative distribution function (CDF) of this same distribution is shown below. **C)** Heatmaps of diffusion coefficients following Bayesian analysis of all unmasked single molecule trajectory data of Halo-TFEB in BeWo cells treated with different drug conditions. Each row corresponds to the distribution of posterior occupations for a single movie file. **D)** The distribution of diffusion coefficient occupancies for all unmasked trajectories of Halo-TFEB treated with DMSO or Forskolin for 48hrs. The fraction bound (calculated by the fraction of the distribution with a diffusion coefficient <0.1$\mu m^2/s$) is shown highlighted with the blue region and the quantified values displayed as black percentages. The cumulative distribution function (CDF) of this same distribution is shown below.

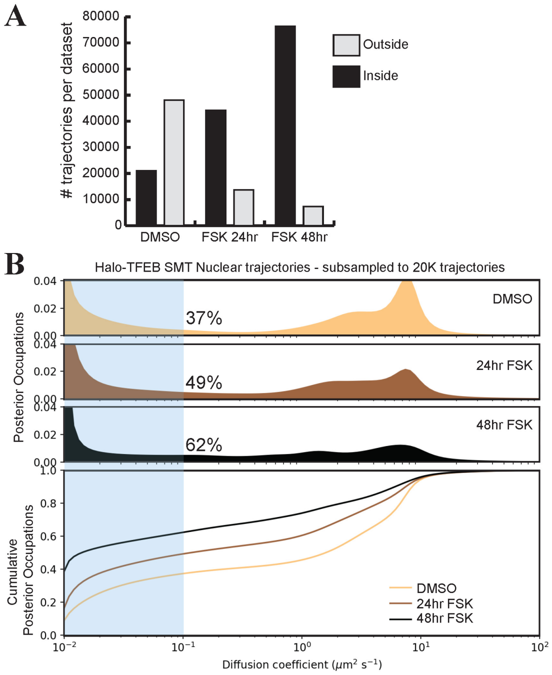


**Figure S5. Subsampling nuclear Halo-TFEB trajectories to assess impact of fewer nuclear trajectories on Bayesian modeling. A)** A barchart of the summed total number of trajectories in each dataset across replicates and cells after masking segmentation. Trajectories fully encompassed inside the nuclear mask (and thus included in Fig 5E) are shown in black bars. **B)** The distribution of diffusion coefficient occupancies after re-analysis of only 20,000 randomly selected masked nuclear trajectories of Halo-TFEB treated in each condition. The fraction bound (calculated by the fraction of the distribution with a diffusion coefficient $<0.1 \mu m^2/s$) is shown highlighted with the blue region and the quantified values displayed as black percentages. The cumulative distribution function (CDF) of this same distribution is shown below.

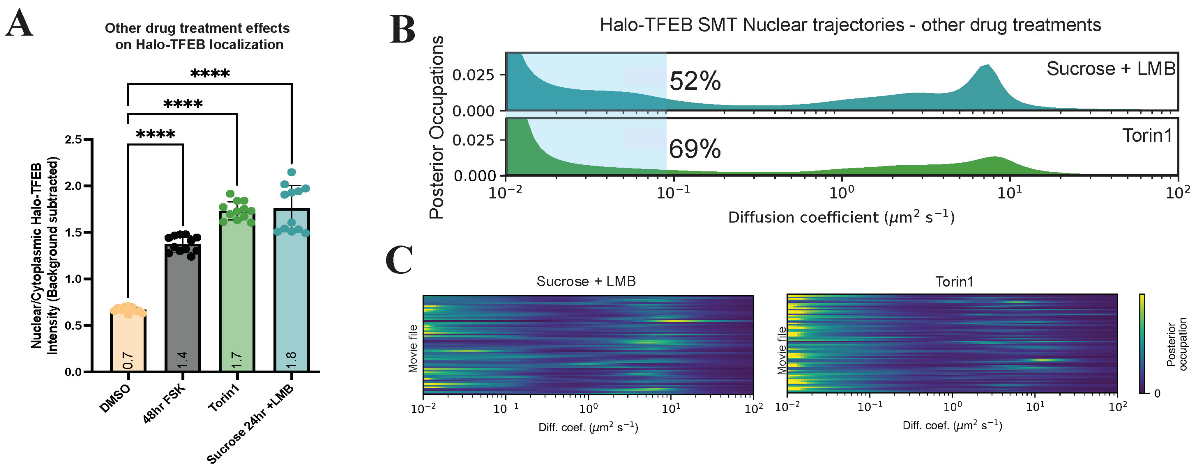


**Figure S6. Nuclear shuttling and chromatin binding analysis of Halo-TFEB upon other drug conditions. A)** Quantification of live cell spinning disk confocal imaging of Halo-TFEB after treatment with specified drug treatment. Statistical significance from an ordinary one-way ANOVA with Tukey's multiple comparisons test is shown where ns = not significant, * = $p<0.05$, ** = $p<0.01$, *** = $p<0.001$ and **** = $p<0.0001$. **B)** The distribution of diffusion coefficient occupancies for nuclear segmented trajectories of Halo-TFEB in other drug conditions. The fraction bound (calculated by the fraction of the distribution with a diffusion coefficient $<0.1\mu m^2/s$) is shown highlighted with the blue region and the quantified values displayed as black percentages. **C)** Heatmaps of diffusion coefficients following Bayesian analysis of single molecule trajectory data of Halo-TFEB in BeWo cells in alternative drug conditions. Each row corresponds to the distribution of posterior occupations for a single movie file.

A

| Replicate | Sample | Overall alignment rate |
|---|---|---|
| Rep 1 | DMSO mFlag | 76.07% |
| Rep 1 | FSK mFLAG | 74.98% |
| Rep 1 | Torin1 mFLAG | 75.95% |
| Rep 1 | DMSO IgG | 89.92% |
| Rep 1 | FSK IgG | 64.55% |
| Rep 1 | Torin1 IgG | 68.79% |
| Rep 1 | DMSO Input | 94.72% |
| Rep 1 | FSK Input | 94.73% |
| Rep 1 | Torin1 Input | 94.36% |
| Rep 2 | DMSO mFlag | 74.56% |
| Rep 2 | FSK mFLAG | 61.96% |
| Rep 2 | Torin1 mFLAG | 67.57% |
| Rep 2 | DMSO IgG | 68.00% |
| Rep 2 | FSK IgG | 69.06% |
| Rep 2 | Torin1 IgG | 70.17% |
| Rep 2 | DMSO Input | 77.71% |
| Rep 2 | FSK Input | 79.01% |
| Rep 2 | Torin1 Input | 76.03% |

B

| Condition | Called against (control sample) | # IDR significant peaks called (p-value<0.05) |
|---|---|---|
| DMSO | IgG | 116 |
| FSK | IgG | 1948 |
| Torin1 | IgG | 3539 |
| ALL | IgG | 5603 |
| DMSO | Input | 197 |
| FSK | Input | 1656 |
| Torin1 | Input | 3393 |
| ALL | Input | 5246 |

C

| Condition | # Peaks differentially enriched (MACS2 bdgdiff) | # diff-enriched peaks called as IDR-significant peaks (p<0.05) | # diff-enriched & IDR-significant peaks mapped to promoters (TSS +/- 2000bp) | # diff-enriched & IDR-significant peaks mapped to distal regions (TSS >2000bp) |
|---|---|---|---|---|
| FSK-enriched | 530 | 175 | 18 | 155 |
| Torin1-enriched | 6312 | 2175 | 444 | 1662 |
| FSK and Torin1 common | 5343 | 2212 | 454 | 1684 |

D

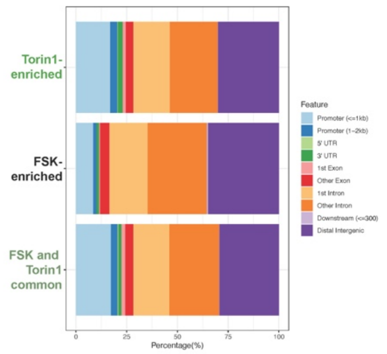


E

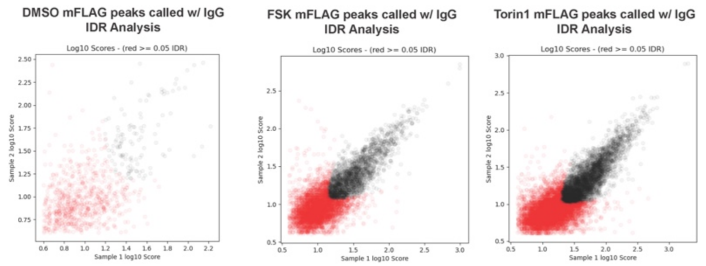


F

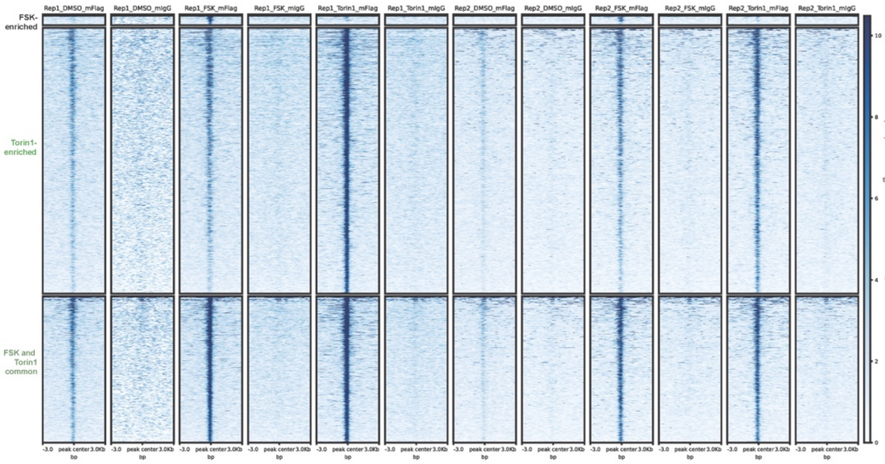


G, H

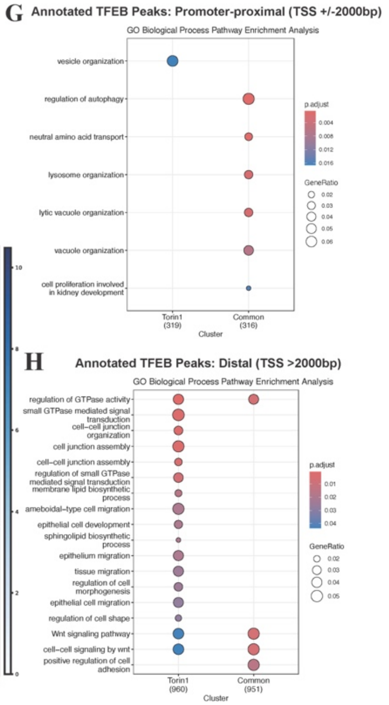


**Fig S7. 3xFlag-Halo-TFEB ChIP quality control and peak filtering. A-C)** Tables indicating the sequencing depths and subsequent filtering steps in ChIP-Seq processing and analysis. **D)** Annotation of IDR robust (score <0.05) peaks to genomic regions using ChIPSeeker. **E)** IDR Analysis of robust peaks across both ChIP-Seq replicates. Sample1 and Sample2 correspond to each replicate and black points indicate robust peaks with an IDR score of <0.05. **F)** Heatmap of differentially enriched peaks analyzed with MACS2 bdgdiff showing +/- 3000bp centered around each peak for all replicates independently. **G)** GO enrichment analysis of all IDR robust (score <0.05) TFEB peaks annotated to promoters (TSS +/- 2000bp) across all samples (Forskolin, Torin1, and DMSO treatments). **H)** GO enrichment analysis of all IDR robust (score <0.05) TFEB peaks not annotated to promoters (distal regions = TSS> 2000bp) across all samples (Forskolin, Torin1, and DMSO treatments).
